# “Online” modulation of brain hemodynamics despite stereotyped running

**DOI:** 10.1101/2020.01.11.902932

**Authors:** Antoine Bergel, Elodie Tiran, Thomas Deffieux, Charlie Demené, Mickaël Tanter, Ivan Cohen

**Affiliations:** Sorbonne Université, CNRS, INSERM, Institut de Biologie Paris Seine-Neuroscience, F-75005 Paris, France; Physique pour la Médecine Paris, INSERM U1273, ESPCI Paris, CNRS FRE 2031, PSL Université Recherche, Paris, France; Université Paris Diderot, Sorbonne Paris Cité, Paris, France

**Author notes:** These authors contributed equally to this work (Mickaël Tanter, Ivan Cohen). Correspondence and request for materials should be addressed to A.B. M.T or I.C.

**Keywords:** Locomotion, Brain hemodynamics, Neurovascular coupling, Ultrasound imaging, Theta rhythm, Gamma rhythms

## Abstract

Theta and gamma rhythms coordinate large cell assemblies during locomotion. Their spread across temporal and spatial scales makes them challenging to observe. Additionally, the metabolic cost of these oscillations and their contribution to neuroimaging signals remains elusive. To finely characterize neurovascular interactions in running rats, we monitored brain hemodynamics with functional ultrasound and hippocampal local field potentials in running rats. Theta rhythm and running speed were strongly coupled to brain hemodynamics in multiple structures, with delays ranging from 0.8 seconds to 1.8 seconds. Surprisingly, hemodynamics was also strongly modulated across trials within the same recording session: cortical hemodynamics sharply decreased after 5-10 runs, while hippocampal hemodynamics strongly and linearly potentiated, particularly in the CA regions. This effect occurred while running speed and theta activity remained constant, and was accompanied by increased power in hippocampal high-frequency oscillations (100-150 Hz). Our findings reveal distinct vascular subnetworks modulated across fast and slow timescales and suggest strong adaptation processes despite stereotyped behavior.

## Introduction

From the early days of electroencephalography (EEG), brain rhythms have been observed in a wide range of models and used as specific markers to characterize specific behaviors such as locomotion, sleep states, attention or cognitive control^1, 2^. Neural oscillations support timely communication between coherent distant brain areas by providing windows of opportunity for efficient spike synchrony^3, 4^ and their disruption often is a hallmark of pathological conditions like epilepsy, schizophrenia, or Parkinson’s disease^5^. Over the past decade, studies have reported that numerous brain rhythms are global processes that are not stationary, but instead circulate across brain regions. During locomotion in rodents, theta waves travel along the septotemporal axis of the hippocampus^6, 7^, slow waves during NREM sleep travel from anterior towards posterior sites^8^ and sleep spindles in humans rotate along a temporal, parietal and frontal cortical sites^9^. Because it is challenging to capture neural activity globally, this poses a problem from an experimental standpoint that high-density recordings can only partially solve, let alone the intrinsic caveats of electrophysiology.

Theta rhythm (6-12 Hz) is extensively studied in behavioral neuroscience, because it is a fundamental model to understand neural synchronization during complex behavior. It is observable in many brain structures (hippocampus, entorhinal cortex, subiculum, striatum and thalamus) and species (bats, cats, rabbits, dogs, rodents, monkeys)^10^ when an animal engages in walking, running, whisking and foraging behaviors or enters rapid-eye-movement sleep^11^. Among the multiple functions attributed to theta rhythm, it is critical for sensorimotor integration^12^, contextual information encoding^13^, hippocampal-cortical communication^14^ and memory consolidation during REM sleep^15^. Most importantly, theta rhythm provides a temporal structure that organizes place cell firing into “theta sequences”, which encode trajectories during active locomotion and are critical for memory consolidation^16^. Another essential feature of theta oscillation is the presence of nested faster (gamma) oscillations in the 30-150 Hz range which exhibit cross-frequency phase-amplitude coupling^17^. In rodents, these oscillations have been divided into three different subtypes, namely low gamma (30-50 Hz), mid gamma (50-100 Hz) and epsilon or high-frequency oscillations (HFO) (100-150 Hz) which are generated by different brain structures^18^ to serve different functions like memory encoding or retrieval^19^.

Besides electrophysiological studies, there is an important knowledge gap about the spatiotemporal dynamics of large-scale brain networks during natural locomotion and their metabolic cost compared to quiet wakefulness. It is reasonable to believe that this cost is elevated due the diversity of brain regions that locomotion requires and the wealth of non-neuronal processes that it triggers including sustained and coordinated muscular activity, elevated heart and respiration rate, increased neuromodulator regulation and astrocytic activity^20, 21^. In addition, the fact that theta rhythm is observable across cerebral structures, suggests that large-scale cell assemblies are active and coordinated during locomotion. To what extent the activity of these cells is reflected in brain hemodynamics and what is the relative cost of different brain rhythms that involve different cell types, are questions of the utmost importance to interpret BOLD-fMRI and hemodynamics signals^22^. The common consensus on neurovascular coupling, is that local neural activation triggers an increase in blood flow to meet higher energy demands and ensure waste products removal^23^. The exact mechanisms of this coupling are complex and involve several pathways in parallel^24^. They show regional-dependence^25^, non-linearity^26^ and cell-type specificity^27^.

To quantify the neurovascular interactions in distributed brain networks (including deep structures) during natural locomotion, we used the recently developed mobile functional ultrasound (mfUS) imaging modality together with extracellular recordings of local field potentials (LFP) in the dorsal hippocampus and video in freely-moving rats. Functional ultrasound can monitor brain hemodynamics over repeated and prolonged periods of time in mobile animals, which makes it a well-suited tool for functional imaging of complex behavior such as locomotion^28^. Its key features include large field of view, high spatial (in-plane:100 μm x 100 μm, out-of-plane: 400 μm) and temporal (200 ms) resolutions and high sensitivity to transient events. We found that locomotion activates a wide network including dorsal hippocampus, dorsal thalamus, temporal and parietal cortices which are activated in a dynamic sequence consistent across recordings. Hippocampal theta and high-gamma rhythms (50-150 Hz) correlated with vascular activity strongly in the hippocampus and thalamus, but only weakly in the neocortex. Intriguingly, brain hemodynamics were strongly modulated across trials and showed a rapid adaptation (amplitude reduction) in cortical regions concurrent with a more gradual linear potentiation in the hippocampus, while running speed and theta rhythm remained constant. We could relate this vascular reshaping to an increase in high-frequency oscillations power within individual recordings. Our data suggest that the stereotyped repetition of locomotion endogenously modulates brain hemodynamics on a rapid (minute) timescale, and does so without noticeable behavioral modification. These results provide new insights into the understanding of neurovascular coupling during locomotion and suggest that two seemingly similar repetition of a given behavior may actually differ totally in terms of brain activity.

## Results

### Voluntary locomotion activates a brain-wide vascular network

To reveal the vascular networks recruited during natural locomotion and finely characterize the neurovascular interactions between hippocampal rhythms and hemodynamics, we used functional ultrasound (mfUS) in mobile rats combined with electrophysiological recordings of local field potentials (LFP) in the dorsal hippocampus and video [Figure 1A], using previously introduced mfUS-LFP-video setup^28^. Animals were trained to run back and forth for water reward on a 2.2-meter linear, before they underwent surgery and were recorded daily after a 7-day recovery period (see Methods). LFP, electromyogram (EMG) and accelerometer (ACC) signals and video were monitored continuously and post-processed to extract animal position, speed, head acceleration and band-specific theta (6-10 Hz), low-gamma (20-50 Hz) mid-gamma (50-100 Hz) and high-gamma power (100-150 Hz). Depending on the setup and apparatus used, functional ultrasound imaging can probe cerebral blood volume (CBV) and cerebral blood flow (CBF) over multiple brain regions, including deep structures, over a single plane or a full volume^29, 30^ with a spatial resolution depending on the frequency of the ultrasonic probe (15 MHz linear probe: 100×100×400 microns resolution) and a temporal resolution up to 500 Hz, when Doppler frames are formed using sliding windows from compound images. In this experiment, we used a ‘burst sequence’ that alternates 12-second sonication periods during which ultrasound echoes are sent and received at 500 Hz until RAM saturates, with refractory periods for ultrasound images to be ‘beamformed’ and transferred, during which vascular activity cannot be monitored. The onset of ultrasound acquisition was triggered manually by the experimenter, when the animal initiated a body rotation to start a new run in the opposite direction on the linear track [Figure 1B]. Such a setup enabled the monitoring of CBV variation during single runs at 100×100×400 µm spatial and 200 milliseconds temporal resolution in multiple brain structures over a single imaging plane. On average, 1 out of 3 runs occurred during the refractory period and could not be imaged, leading to an average of 30-35 trials over 100 runs per recording, with a global balance between each running direction. The tracking of the animal’s position enabled to realign all 12-second fUS recording periods towards a common time origin, defined as the onset of each run [Figure 1C]. A single trial was defined as a run that was correctly captured from onset to end in the 12-second sequence, with run onset and end defined as a 10%-threshold of peak speed (see Methods).

**Figure 1:**
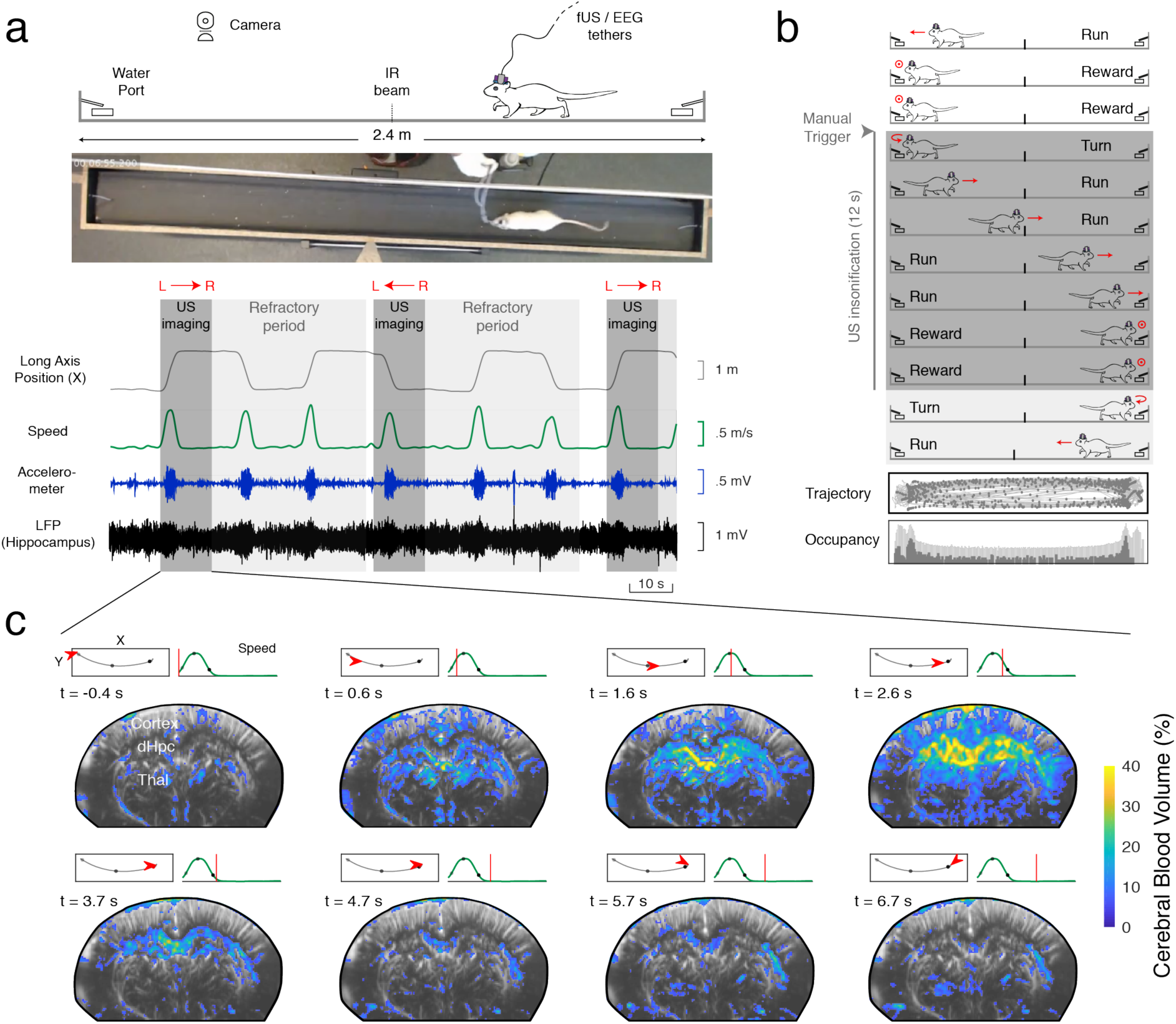
Simultaneous electrophysiological/neurofunctional ultrasound recording reveals single-run hemodynamics in deep brain structures. (a) Schematic and picture of the mfUS-LFP-VIDEO setup. Rats are trained daily to run back and forth on a 2.2 m linear track for water reward, given at both ends. Video, electrophysiology and accelerometer data are acquired continuously. Ultrasound data is acquired in bouts lasting 12 seconds (shaded dark grey), interleaved with a 40 second refractory period (shaded light grey), during which ultrasound data is beamformed. Animal position on the track (grey) and speed (dark green) are extracted offline concurrently with LFP processing. (b) Details of single-run imaging. Ultrasound acquisition is triggered manually by the experimenter, when the animal initiates its turn. A typical 12-s imaging sequence contains Turn (1s) – Run (3-4s) – Reward (7-8 s) recording frames. A typical recording session lasts 30 to 40 minutes and contains 80-100 trials, a third of which are imaged (2/3 occurring during the refractory period). Trajectory and occupancy of the track are displayed at the bottom for the full session (gray) and for the imaged runs (blue dot: one Doppler frame, 200 milliseconds compounding). (c) Spatio-temporal dynamics of the vascular system during a single run. A typical coronal plane (Bregma −4.0 mm) reveals cerebral blood volume (CBV) in cortical, hippocampal and thalamic regions. For each 12 s US insonification, we formed 60 Power Doppler frames (200 milliseconds compounding, 8 frames showed, 5 Hz sampling) based on ultrasound echoes. This reveals prominent activation in the dorsal hippocampus (30-40 %), peaking 2.5 s after US onset, that is 1.5 to 2.0 s after run onset. Timing from run onset is given for each frame. Top left: Animal position on the track (red arrow) overlaid on trajectory (black) Top right: Animal speed as a function of time (red line: current position). Grey dots mark run onset (light grey), run peak (grey), and run end (dark grey).

To reveal the hemodynamic responses to locomotion in a large number of regions while keeping sufficient statistical power, we chose to perform repeated ultrasound acquisitions in different animals over two typical recording planes: a coronal plane (antero-posterior axis, AP = −4.0 mm) that intersected dorsal hippocampus, thalamus, hypothalamus and cortex, including auditory (AC), primary somatosensory barrel field (S1BF) lateral parietal association (LPtA) and retrosplenial (RS) cortices, and a diagonal plane tilted 45° relative to coronal view that intersected hippocampus (dorsal, intermediate and ventral), thalamus (dorsal and ventral), cortex (anterior, midline and posterior) and caudate Putamen (CPu) [Figure 2A – Figure S1]. We then registered two-dimensional vascular planes over reference atlases (Paxinos atlas for coronal planes; Waxholm MRI atlas for diagonal planes) to derive regions of interests (ROIs) as described previously^31^ and spatially averaged the CBV signal on these regions of interest (ROIs). We re-aligned all trials for all recordings to the onset of each run and computed temporal averages of brain hemodynamics of all aligned trials and compared them with behavioral parameters such as running speed and head acceleration [Figure 2B]. This revealed prominent CBV increases that were time-locked to run onset in the dorsal hippocampus bilaterally though both inter-trial and inter-individual variability were present. To characterize the vascular network involved in locomotion we displayed the average hemodynamic responses for 16 brain regions identifiable on most coronal recordings (11 animals, 22 recordings, 384 trials) together with the average running speed. This revealed prominent activations in lateral parietal and retrosplenial cortices, all dorsal hippocampal sub-regions with strongest activation found in the dentate gyrus, and dorsal thalamus [Figure 2C]. A similar analysis over diagonal plane recordings revealed that parietal cortical regions and posterior cortical regions were also active during locomotion, that dorsal hippocampus was strongly recruited while ventral hippocampal activations were absent, and that such dissociation was also visible in the thalamus [Figure S1]. Interestingly, hemodynamic responses were very symmetrical and showed no dependence on running direction [Figure S2].

**Figure 2:**
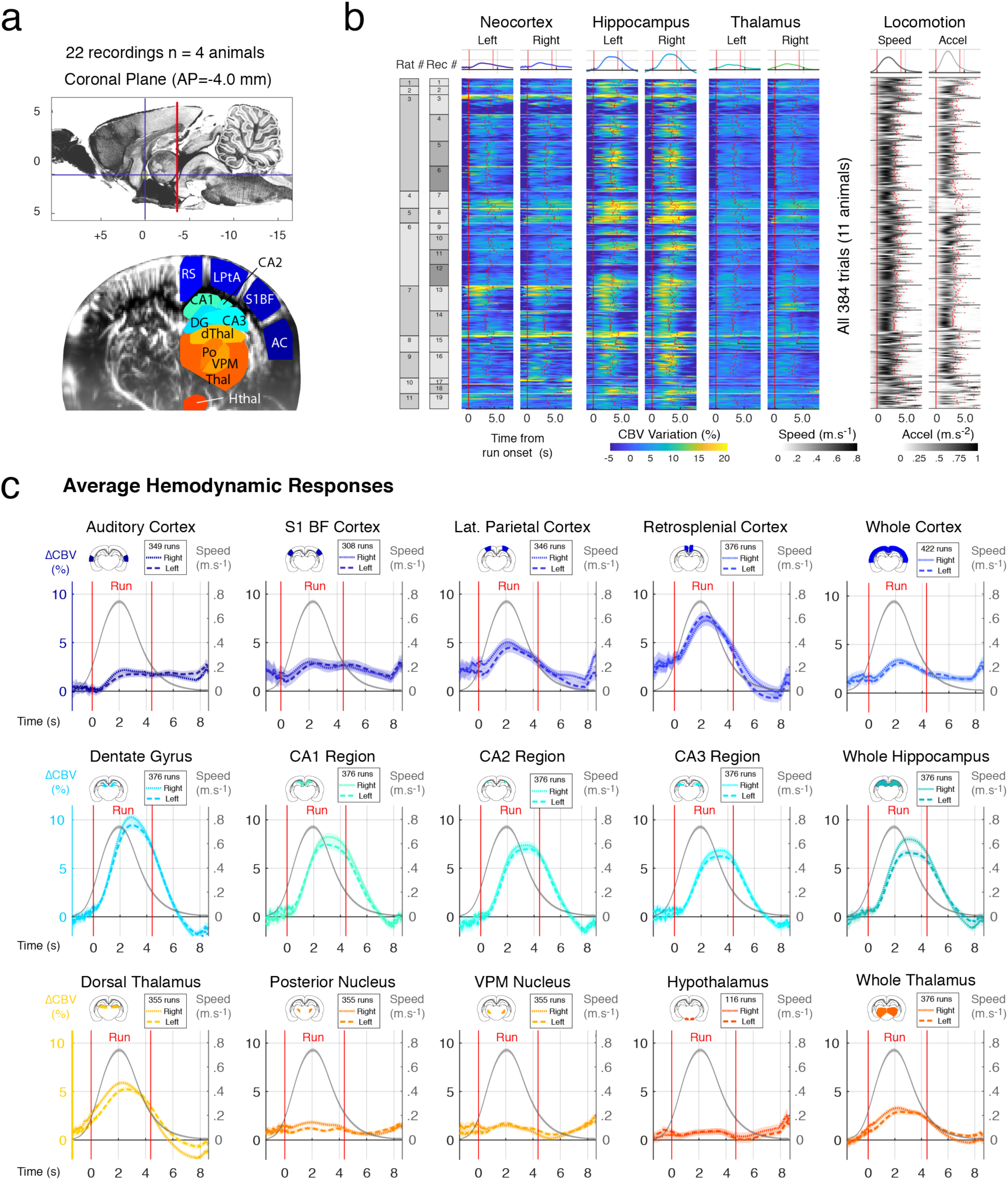
Large-scale hemodynamic responses to locomotion in freely-running rats. (a) Location of a typical coronal (Bregma = −4.0 mm) recording plane and associated brain structures monitored during mfUS-EEG recordings. Top: Sagittal view of the rat brain of the imaging plane position (red) on atlas background. Bottom: Power Doppler image before (left) and after (right) atlas registration, performed by positioning salient landmarks onto Power Doppler image and registering a 3D volumetric segmented MRI atlas. RS: retrosplenial cortex, LPtA: lateral parietal associative cortex, S1BF: S1 barrel field, AC: auditory cortex, DG: dentate gyrus, CA1-CA2-CA3 region, dThal: dorsal Thalamus, Po: Posterior Thalamic Nucleus, VPM: Ventroposterior Thalamic Nucleus, Thal: Thalamus, Hthal: Hypothalamus. (b) Bulk representation of single-trial hemodynamic responses to locomotion in six major brain regions (Neocortex, Hippocampus, Thalamus, left and right). A total of 384 trials (19 recordings, 7 animals) was acquired across many days. For each run, the onset of movement is used as a temporal reference (zero-timing) and all trials are aligned to run onset (see Methods). Run end however is different for all trials. We can then compute average hemodynamic responses (top) and average run duration (t = 4.26s +/- 0.3s). The same approach is performed for locomotion parameters: running speed (left) and head acceleration (right). Hemodynamic responses are symmetrical and display strong inter-trial variability in the cortical and hippocampal regions, with strongest activations in the dorsal hippocampus. (c) Average hemodynamics responses over multiple regions. Using the approach in b, we computed average responses to locomotion in 24 regions. We overlay running speed (grey) to visualize both the response intensity and temporal delays to locomotion. Overall, strong bilateral activations are found in all dorsal hippocampus regions (stronger and earlier in the dentate gyrus), retrosplenial cortex and dorsal thalamus. We display hemodynamic responses for large regions as a reference on the right side. For each region the number of runs can differ slightly as all 24 regions were not always visible on each recording.

### Distant brain regions are activated in a dynamic sequence with delays ranging from 0.8 to 1.8 seconds after peak running speed

In order to quantify the coupling between brain hemodynamics and running speed in different brain and to reconstruct the sequence of activation during voluntary locomotion, we computed cross-correlation functions between CBV pixel signals and running speed [Figure 3A]. Importantly, cross-correlation is not precluded by the discontinuity of CBV signals (due to the refractory period imposed by the burst sequence) that display long periods without signal, because speed was monitored continuously meaning that CBV-speed correlations can be computed for all time delays, in a similar fashion as what would be done if CBV signal were continuous. For all pixels in a given recording, we extracted the coordinates of the peak of the cross-correlation function leading to a measure of maximal correlation R_max_ and a corresponding delay T_max_. We then displayed these cross-correlation functions grouping them by anatomical ROIs as defined in Fig. 2. Strong cross-correlation values were prominent in the dorsal hippocampus, dorsal thalamus and in parietal and posterior cortices. We also reconstructed peak correlation and peak time maps to represent visually which regions displayed the strongest correlations and their corresponding timing [Figure 3B]. These maps were consistent with previous results, showing strong correlation coefficients in the dorsal hippocampus and dorsal thalamus bilaterally. They also revealed that variability was stronger in cortical regions and that significant time delays were visible between the dorsal and ventral parts of the thalamus and between anterior and posterior structures in diagonal recordings. This approach led to a precise time sequence of individual pixel responses to locomotion, but did not allow for statistical comparison across recordings, due to pixel mismatch resulting from heterogenous probe position across sessions.

**Figure 3:**
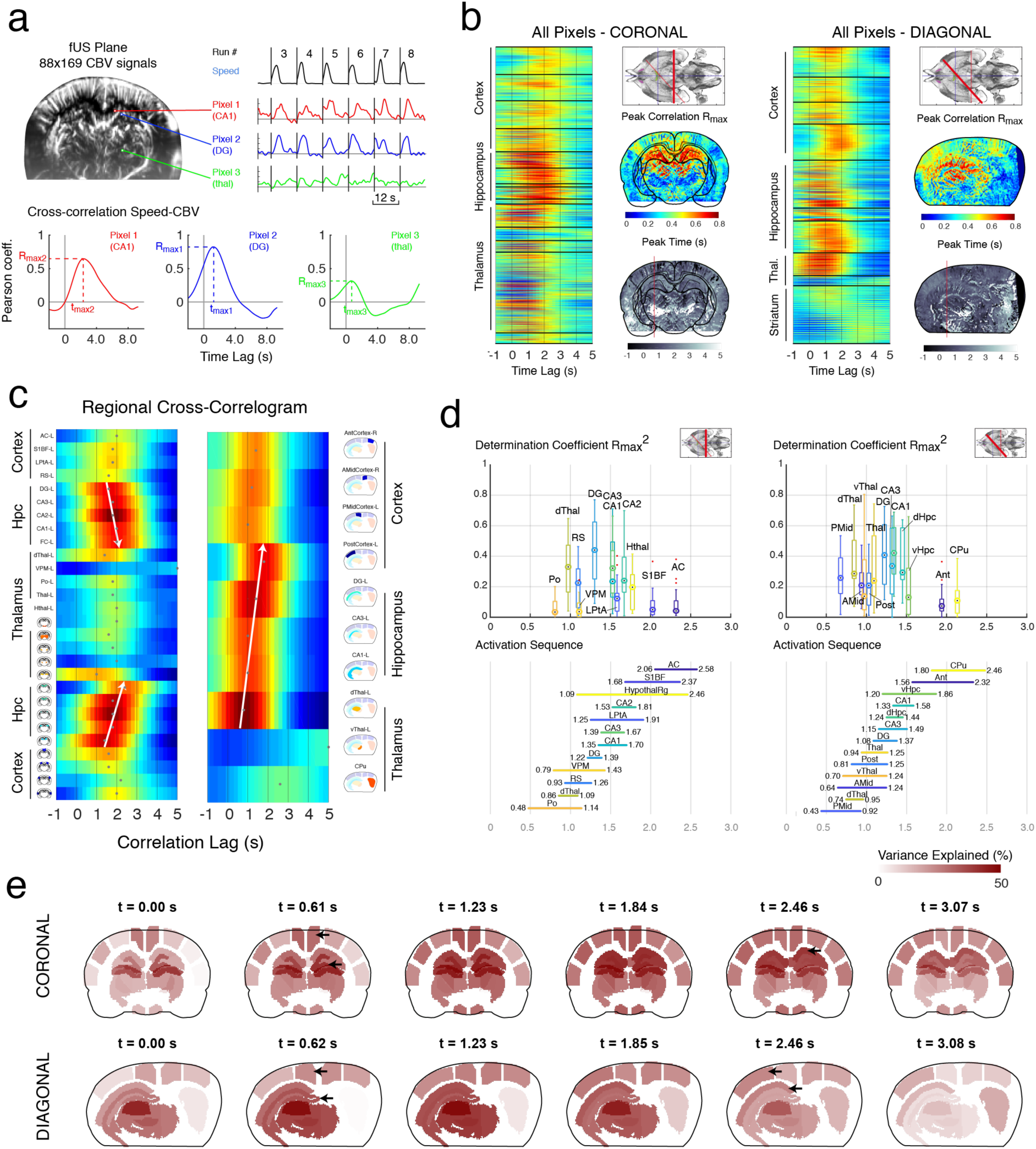
The vascular network responds dynamically to locomotion with sequential propagation along the tri-synaptic loop at the sub-second timescale. (a) fUS data is acquired along a typical coronal plane (Bregma = −4.0 mm, N=22 recordings, 7 animals) and a diagonal plane (Delta = 45-60°, N=20 recordings, 7 animals). A typical plane contains 15,000 voxels from which a CBV signal can be extracted and synchronized with speed (Top right, example of 3 pixels, red CA1 region, blue Dentate gyrus, green Po nucleus). For each pixel, we compute the cross-correlation function (time-lags from −1.0 s to 5.0 s) between speed and CBV signal. Then we extract the peak coordinates, leading to a measure of the peak correlation R_max_ and corresponding peak time T_max_. (b) The approach illustrated in (a) is performed for all pixels in two typical coronal (left) and diagonal (right) recordings. After atlas registration, we can re-order the cross-correlations by regions. Strong correlations are clearly visible is some cortical regions (RS) and thalamic (dorsal) regions, while all dorsal hippocampal regions strongly correlate with speed. Note the different delays across brain regions. For each recording, we can extract a peak correlation map (top) and a peak timing map (bottom). Note the consistency between the two recordings taken from 2 different animals. (c) Regional correlograms for the two recordings shown in (b). We derived an average CBV signal across each region and computed its cross-correlation with speed, similarly to (b). This reveals a dynamic pattern of activation across regions at the sub-second timescale. Note the clear propagation along the regions of the dorsal hippocampus bilaterally (white arrows). This pattern is visible but less clear on the diagonal plane, due to a more challenging registration. Ventral thalamus, striatum and hypothalamus respond weakly to locomotion. For each region in each recording, we can extract a peak correlation R_max_ and corresponding peak time T_max_, to be compared across individuals. (d) Synthesis of determination coefficients (R_max_^2^) and time delays for all coronal and diagonal recordings. Note that the strongly responsive regions shown in Figure 2 (retrosplenial cortex, dorsal thalamus, dorsal hippocampus) strongly correlate with animal speed. Dentate gyrus and CA regions show a consistent activation delay (300-500 milliseconds). The discrepancies between coronal and diagonal planes are probably due to the inhomogeneities in spatial averaging over AP axis in diagonal planes. Interestingly, ventral thalamus and striatal regions are silent during locomotion and active after run end. (e) Post-hoc reconstruction of the locomotion-related vascular propagation based on the mean peak correlation and peak time for each region, computed in (d). The transparency of each region is proportional to the Pearson correlation coefficient at the time-lag t specified on the top of the graph. The sequential tri-synaptic wave is visible on the reconstruction. Note that the parietal cortex is concurrent with dentate activation while posterior cortex co-occurs with downstream CA1 activation.

To circumvent this and reconstruct a dynamic sequence across individuals, we analyzed the speed-CBV cross-correlations of regional hemodynamics. In coronal observations [Figure 3C & 3D], we observed that dorsal thalamus (0.98 s ± .12 s) and retrosplenial cortex (1.03 s ± .13) peaked earlier than hippocampal regions. When focusing on hippocampal sub-regions, we found that dentate gyrus peaked earlier (1.31 s ± .09 s, 15 sessions, n = 4 animals) than CA3 (1.58 s ± .09 s) CA1 (1.58 s ± .12 s) and CA2 (1.69 s ± .11 s) regions. This pattern was bilateral and robust across recording sessions. Average dentate gyrus response typically started earlier and lasted longer than CA1/CA2/CA3 responses, which were shorter, but of higher amplitude. This suggests that hippocampal subfields are perfused by at least two distinct vascular networks. Consistent with earlier observations, ventral thalamus displayed moderate anti-correlation relative to animal speed^28^. The dynamic sequence of vascular activation was also visible over diagonal planes [Figure 3D] starting in the parietal cortex (1.31 s ± .09 s) dorsal thalamus (1.31 s ± .09 s), then reaching dentate gyrus (1.31 s ± .09 s) and CA1 region (1.31 s ± .09 s). Interestingly, relative delays across regions were consistent across coronal and diagonal planes but absolute speed-CBV delays tended to be shorter in diagonal planes. This can be explained by the fact that accurate atlas registration was more error-prone on these recordings, especially in hippocampal subfields but also by the fact that diagonal plane recordings included more anterior regions that could potentially peak earlier than temporal sites, thus biasing the average correlation delay towards earlier values. Our results suggest a pattern of activation in the dorsal two-thirds of the hippocampus. The medial septum was visible on some recordings but not clearly identifiable across recordings which impeded statistical assessment. On some recordings, hemodynamics in the subiculum could be observed, revealing a continuation of the delayed activation in CA1 (R_max_ = 0.39 ± .15, T_max_ = 2.46 ± .95, 8 sessions, n = 2 animals, not shown).

In order to exhibit the fine spatiotemporal dynamics of the speed-CBV correlations, we reconstructed high-definition Doppler movies using sliding 200-ms sliding windows, with 190-ms overlap from underlying 500 Hz ultrasound data^32^. This revealed the fine dynamics of brain hemodynamics at very high temporal resolution (10 ms) in two typical coronal and diagonal recordings [Movie S1-S2]. Ultimately, we reconstructed a ‘reference’ activation sequence based on the average measures (R_max_,T_max_) for each ROI from all individuals [Figure 3E] to give an overview of the spatiotemporal dynamics of the sequence associated with running speed over the two chosen recordings planes.

Theta, mid gamma and high bursts of activity were clearly visible on the spectrogram when the animal prepared to or actually engaged in running on the linear track, but not during reward uptake [Figure 4A]. Low-gamma and high-gamma oscillations were nested in theta oscillations and occurred slightly later, with low gamma episodes occurring less frequently, consistent with the involvement of low gamma in memory retrieval and that of high gamma in memory encoding^19^. Theta and gamma showed phase-amplitude cross-frequency coupling, with low-gamma peaking on the ascending phase of theta, mid-gamma power being maximal at theta peak and high-gamma maximal at theta trough [Figure 4B]. This LFP signature of CA1 recordings is in accordance with previous results in the literature^18, 33^. To go one step further and investigate the nature of neurovascular interactions between hippocampal rhythms and CBV signals, we computed LFP-CBV cross-correlation functions between four regional signals (cortex, whole hippocampus, whole thalamus and whole-brain) and four typical LFP bands: theta (6-10 Hz), low gamma (20-50 Hz), mid gamma (50-100 Hz) and high gamma (100-150 Hz) [Figure 4C]. This analysis has been performed previously during REM sleep^31^ and showed high-gamma oscillations strongly correlated with vascular activity in most brain regions (theta oscillations also, but more moderately). During locomotion however, theta rhythm correlated with vascular activity more strongly than mid and high gamma in all brain regions. Additionally, neurovascular coupling shows a stronger regional dependence: whereas REM-sleep hemodynamics activations were hypersynchronous across brain regions, revealing very strong LFP-CBV coupling in most regions, locomotion-related hemodynamics were restricted to a subset of precise (but distributed) regions as detailed earlier. Hence, hippocampal theta activity was more strongly associated with vascular responses in the hippocampus and thalamus (R_mean_ = 0.51, R_mean_ = 0.40), and markedly less in cortical regions (R_mean_ = 0.28). Consistently with previous results, low gamma did not correlate with vascular responses in any brain region, and mid and high-gamma bands showed moderate correlation with CBV responses only in the hippocampus (R_max_ = 0.41, R_max_ = 0.41 in both regions). The analysis of the distributions of LFP-CBV delays (direct measure of neurovascular coupling delays) revealed that LFP always preceded vascular activity [Figure 4D]. The earliest delays were consistently found in the thalamus (1.32 ± .08 s from theta peak) followed by the hippocampus (1.71 ± .07 s) and cortex (2.03 ± .12 s). Delay distributions were more homogenous in the hippocampus and thalamus both for theta and gamma bands, meaning that neurovascular coupling was stronger during locomotion in subcortical structures compared to cortical ones. A more detailed analysis of LFP-CBV delays and coupling strength across regions was performed across all coronal and diagonal plane recordings [Figure S3 & S4]. Taken together, these results show that theta activity is a better regressor to explain hemodynamics variance during locomotion and that the robustness of this correlation is stronger in hippocampal and thalamic regions.

**Figure 4:**
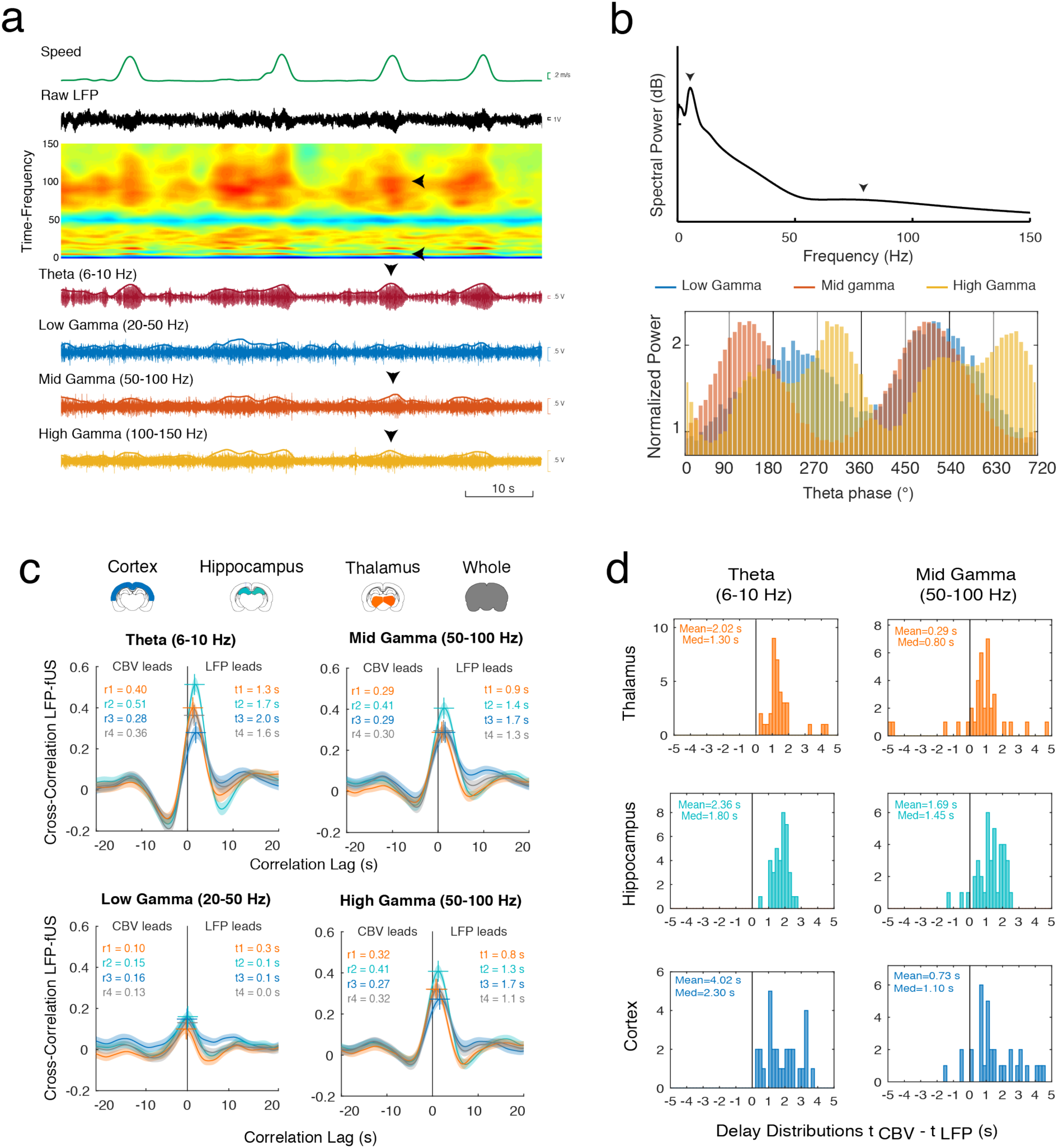
Theta (6-10 Hz) and Fast Gamma (50-150 Hz) precede and correlate with hemodynamic signals during locomotion. (a) Time-frequency analysis of the hippocampal LFP during running. (Top to bottom) Animal Speed, Raw LFP, Time-frequency spectrogram, Bandpass-filtered LFP and power envelopes in the theta (6-10 Hz), low gamma (20-50 Hz), mid gamma (50-100 Hz), high gamma (100-150 Hz) bands. Instantaneous power envelope is computed by temporal smoothing (gaussian kernel 0.5 s). Hemodynamic signals. Note the prominent bursts of theta and fast gamma oscillations concurrent with each individual run (black arrows). (b) Top: Power spectral density of the hippocampal LFP recording during locomotion. Note the two prominent peaks (arrows) in the theta and gamma bands. Bottom: Phase-amplitude cross-frequency coupling between theta and gamma bands in the hippocampal LFP show three distinct sub-bands (0°: theta trough). Theta-phase is extracted by linear interpolation of the zero-crossing function of theta-filtered LFP signal. Low gamma is maximal around at the ascending phase of theta [180°-270°]. Mid gamma is maximal at the peak [90°-180°] and high gamma is maximal close to the trough [300°-360°]. (c) Cross-correlation functions between the four LFP envelope signals shown in (a) the four CBV signals extracted from regions shown on top (Thalamus, Hippocampus, Cortex, Whole). The cross-correlation functions are obtained by averaging the cross-correlation for each recording (N=42 recordings, 7 rats). The x-coordinate of the maximum of this function (color crosses) gives the delay between LFP and CBV signals, while the y-coordinate gives the strength of the coupling. Low-gamma shows no coupling with CBV signals in either region, both mid and high-gamma band show moderate coupling, and theta band shows strong robust coupling especially in the dorsal hippocampus. Note the consistent shorter delays in the thalamus, hippocampus and cortex, confirming the sequential activation of these three structures, shown in Figure 3. Shaded patches show the standard error of the mean. (d) Histogram counts of LFP-CBV delays extracted from the peak of cross-correlation function for two typical bands (theta and mid gamma) and three regions (cortex, thalamus, hippocampus). None of the recordings displayed negative delays, meaning that hemodynamics always follows LFP activity. Variability is stronger in the 1–3 s lag for the cortical responses than for thalamic and hippocampal responses, showing that the thalamic and hippocampal hemodynamic responses are strongly coupled to hippocampal local field potentials, but partially decoupled in the cortical regions.

### Locomotion-related hemodynamics are strongly modulated despite stereotyped running

Because of its high sensitivity and high temporal resolution, transient events can be monitored with fUS circumventing the need for temporal averaging routinely implemented in BOLD-fMRI or PET studies. We thus interrogated whether brain hemodynamics were invariant or modulated across trials within a single recording session and, if such effect was present, whether it affected brain regions to the same extent. We repeated the approach from Fig.2 that consisted in re-aligning all trials onto a common time origin taken as the onset of each run, but this time we focused on each individual recording independently. Brain hemodynamics showed strong modulation across trials within the same recording session, while behavioral parameters such as running speed or head acceleration remained constant [Figure 5A]. We represented hemodynamic responses to individual trials in six locomotion-sensitive brain regions found previously (retrosplenial cortex, dentate gyrus, CA1, CA2, CA3 and dorsal thalamus) by color-coding their amplitude of response, normalized to the weakest and strongest responses within the session (blue: strong response, light green: weak response). While body speed or dorsal thalamus hemodynamics displayed an even repartition of weak and strong responses along the whole session, hemodynamics in the retrosplenial cortex on the one hand and in the four sub-regions of the hippocampus on the other hand showed a strong bias in this repartition: strong cortical activations occurred consistently in the early runs while strong hippocampal trials were gathered in the last 2/3 of the session. This effect was even stronger in the CA regions compared to dentate gyrus. To visualize this finding, three typical runs taken during the early, intermediate or late third of the session were aligned and displayed in [Figure 5B]. In the early run, locomotion strongly activated various cortical regions including retrosplenial cortex together with dorsal thalamus and dentate gyrus. In the intermediate run, the cortical activation was weakened and the hippocampal activation spread to the CA1/CA2/CA3 regions. In the late run, cortical response was absent, thalamic activation was weakened, while the full dorsal hippocampus was recruited massively. The full sequence of individual runs for 2 typical recordings is given in supplementary information [Movie S3-S4].

**Figure 5:**
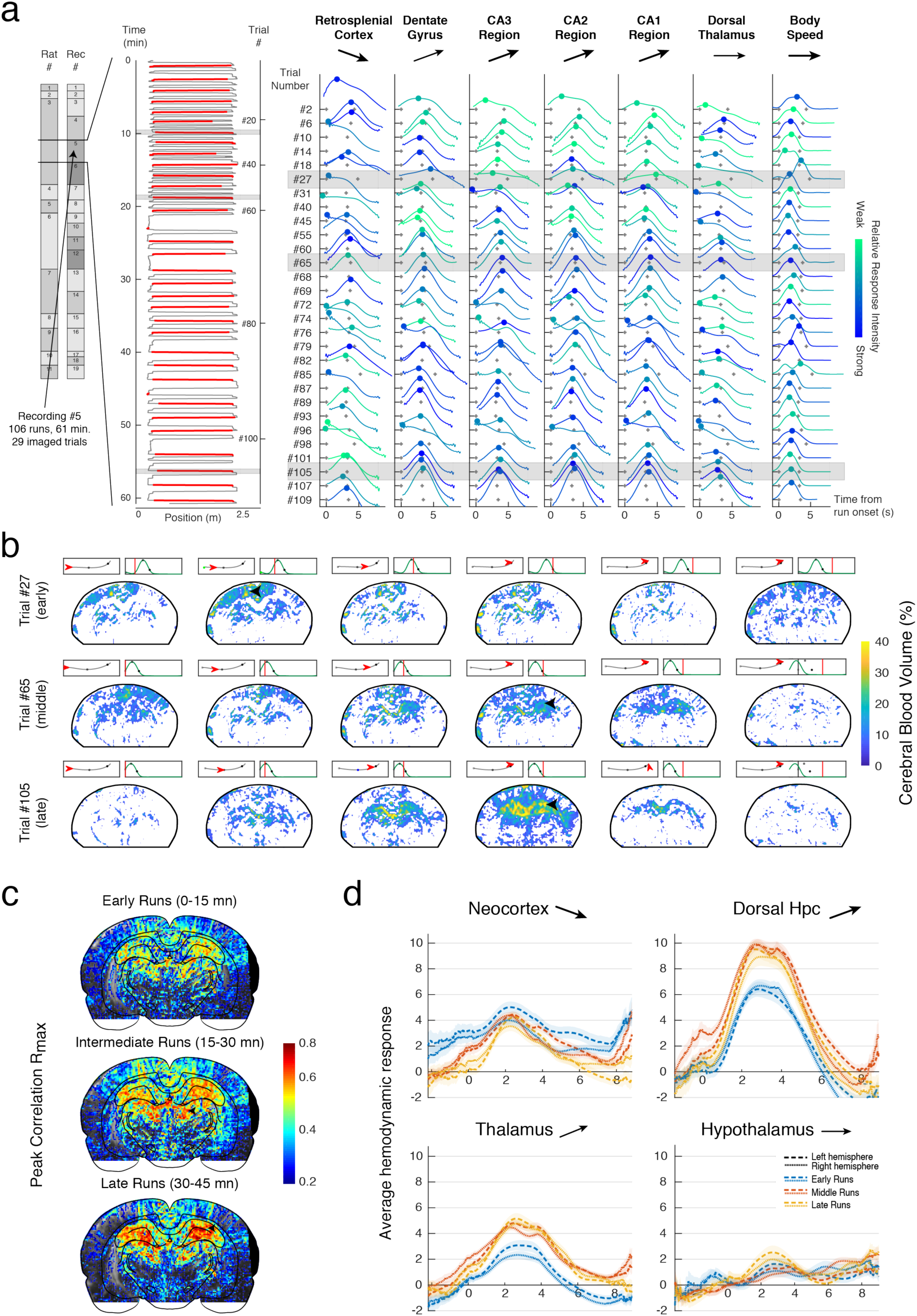
Strong region-dependent plasticity in the vascular network over the timescale of minutes. (a) Inter-trial analysis of locomotion-related hemodynamics is shown for a single recording session, totalizing 106 runs, 29 of which have been captured by fUS. Left: Animal position on the long axis of the linear track versus time. Recorded runs are shown in red (Burst mode imposes a dead time between two 12 s insonification epochs). Right: Regional hemodynamic responses during all 29 imaged trials for the six strongly responsive regions shown in Fig. 2. All six brain regions display inter-trial variability while running speed remained constant (right). For each column, trials are color-coded with respect to their strength of response over the full recording (weak: light green, strong: blue). Black arrows indicate the tendency for the vascular response to either potentiate, remain stable or decrease (bold: strong tendency). Hippocampal regions potentiate very strongly across trials, while retrosplenial cortex depresses. Dorsal thalamus and dentate gyrus to a lesser extent remained relatively constant. Importantly, speed remains relatively constant throughout the session. (b) Spatiotemporal dynamics of 3 trials shaded in (a), each taken during the early (trial 6), middle (trial 12) and last (trial 27) portion of the recording. For the early trial, we observe widespread cortical activation and moderate dentate and dorsal thalamus activation. For the intermediate, the hemodynamic response spreads to the downstream regions of the hippocampus (black arrow). For the late trial, cortical response is absent while the hippocampus responds very strongly at the end of the trial (black arrow). (c) Maximal correlation maps relative to animal speed, as shown in Fig. 3 for three trial batches grouping early (0-15 min from recording onset), intermediate (15-30 min) and late (>30 min) runs. The strong potentiation in the downstream regions of the dorsal hippocampus shows that locomotion-related hemodynamic responses shift from cortico-thalamic sites to regions downstream of the hippocampus. (d) Average hemodynamic responses in 4 major brain regions depending on trial timing, grouped by early (0-15 min), intermediate (15-30 min) and late (>30 min) trial groups. Neocortex and hippocampus/thalamus regions display opposing plasticity patterns, showing that potentiation observed through fUS is not a global effect of temperature or reward during the task. Arrows indicate vascular tendency as in (a).

For each recording, we then sorted all trials in three different time groups, respectively including all early trials (0-15 min from session onset), intermediate trials (15-30 min) and late trials (>30 min) in order to compare brain activity across regions and time. Individual pixel-based peak correlation maps showed an increased speed-CBV coupling in the dorsal hippocampus in the intermediate and late groups versus the early time group, concurrent with a decreased speed-CBV coupling in the cortex and in the dorsal thalamus, though the second was more gradual than the first [Figure 5C]. This effect was consistent across recordings: the average hemodynamic responses for all coronal recordings showed a strong amplitude increase in intermediate/late time groups compared to the early one both in the thalamus and hippocampus, a moderate amplitude decrease in the cortex, while the hypothalamic response remained constant across time groups [Figure 5D]. This effect was also observable in the bulk representations of single-trial responses across regions [Figure S5] Importantly, hippocampal/thalamic hemodynamics on the one hand and cortical hemodynamics on the other, were strongly reshaped across trials but followed opposite patterns. The fact that some regions showed absent modulation rules out any global effect affecting all regions simultaneously but advocates instead for a strong region-dependent tightly-regulated mechanism yet to be defined.

To assess the temporal dynamics of this vascular reshaping across brain regions, we divided recordings into 1-minute duration bins and computed the median value of regional CBV signals for all trials. We then derived a mean CBV value per bin for all regions across recordings [Figure 6A]. We observed a strong linear progression in CA1, CA2, CA3 regions between peak amplitude response and session onset. Comparatively, dentate gyrus showed slightly lower increase as the peak response in early trials was already significant. In comparison retrosplenial cortex and somatosensory cortex vascular responses were consistently larger in the very early trials (1-5 mins) than for the rest of the task. Dorsal thalamus also displayed a linear increase in peak response that reach a plateau after 10 minutes. Ventral thalamic regions showed relative low dependence on trial timing. We also computed other parameters (CBV value at trial start, trial end, mean across trial) and demonstrated that hemodynamic reshaping mainly affects the late component of the hemodynamic response to single trials. Linear regression of this histograms allowed us to group brain regions into 3 main clusters according to their modulation profile [Figure 6B]. Cluster 1 contained cortical regions that display strong early responses, and moderate depression across trials. Cluster 2 includes regions like the thalamus which display moderate responses in early trials and moderate potentiation. Cluster 3 gathers hippocampal subfields which show low to absent early responses and moderate to strong linear potentiation. In individual recordings, we could then perform linear regression for all pixels and segregate them according to the sign of the potentiation slope. In some recordings, vascular depression was observable in the dentate gyrus and cortical sites, while potentiation was present in the CA1/CA2/CA3 regions of the hippocampus [Figure 6C]. This shows that vascular reshaping affects differentially hippocampal subfields at least in some individuals. Thus, the function of hemodynamic modulation must to some extent relate to functional or anatomical differences between hippocampal subfields.

**Figure 6:**
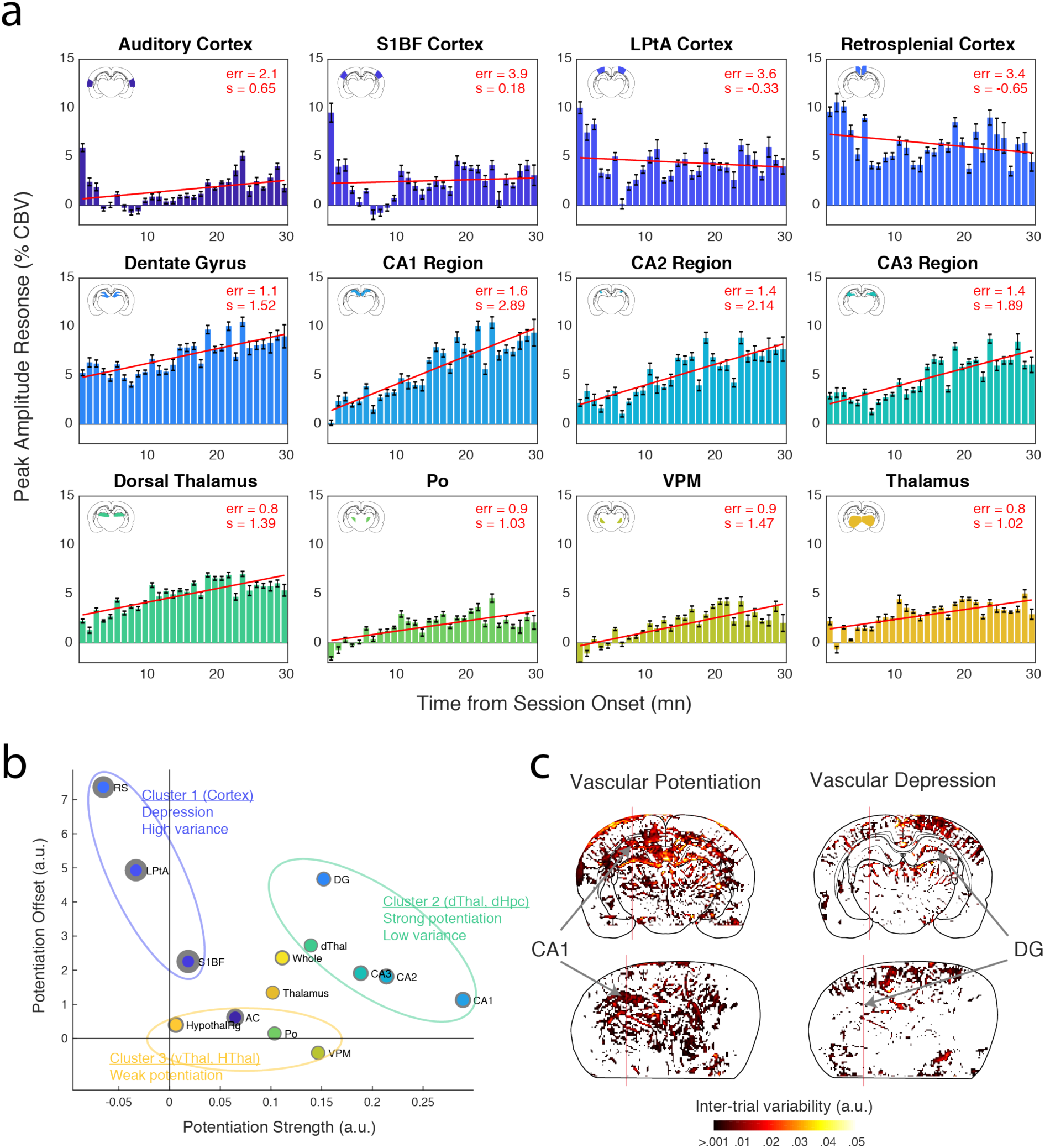
Fine temporal dynamics of vascular plasticity across brain regions. (a) Average peak CBV response to locomotion grouped by timing from session start (0 refers to the first run in each recording) for 12 regions across individuals (N= 7 animals, 22 recordings). Left columns represent early runs while right columns represent late runs. Linear regression lines are shown in red (top corners: err - average quadratic error and s - regression slope). Note that cortical responses are highly variable and show a marked decrease after the first 3-5 min. Hippocampal regions and dorsal thalamus however display a linear dependence on trial timing, with the strongest increase found in the CA1 region. Red lines show the linear regression for left and right regions (b) Clustering of brain regions relative to their vascular potentiation profile. Linear regression-derived potentiation strength (slope in (a)) versus potentiation offset (y-intercept in (a)) for all 12 brain regions. Gray circles lines represent 95% confidence intervals for these 2 parameters. Three main clusters appear in this representation: Cortical regions show a weak linear dependence on trial time, with a marked decrease after early trials and re-potentiation after 10 min (Cluster 1). Dorsal hippocampus and dorsal thalamus display a linear increase in vascular response from early to left trials, with a strongest effect in the CA regions of the hippocampus (Cluster 2). Ventral thalamus, auditory cortex and hypothalamus show weak to absent vascular potentiation (Cluster 3). Grey borders are proportional to the global error of the linear fit. (c) Spatial maps of vascular potentiation profile for all pixels in two typical recordings. The color-code is proportional to potentiation strength (slope in (a)) for all CBV pixel signals in a coronal (top) and a diagonal recording (bottom). Positive (left) and negative (right) potentiation slopes are displayed onto two different graphs. Note the difference between cortical regions on the one hand and dorsal hippocampal regions on the other hand, illustrating the two first clusters in (b).

### High-frequency oscillations but not theta rhythm nor behavioral parameters account for hemodynamic reshaping

Up to this point, we demonstrated that brain hemodynamics strongly reshaped across individual trials within a single recording session; we quantified the precise dynamics of this modulation and showed that this occurred independently of running speed. We then questioned whether vascular reshaping could be accounted for by behavioral parameters or electrophysiological activity, by displaying in parallel vascular, behavioral and electrophysiological signal in each recording. As shown before, hemodynamics showed different modulation patterns in retrosplenial cortex, CA1 region and dentate gyrus while speed and head acceleration remained constant or slightly decreased. Interestingly, we found that while theta activity and low gamma activity remained constant across trials, mid gamma (50-100 Hz) and high frequency oscillations (100-150 Hz) were also positively modulated across trials [Figure 7A]. To provide a quantitative measure of this effect, we computed a median value in each trial for all these variables. This led to column vector of median values, the size of which was the number of trials. We then computed correlations between all possible behavior/CBV and electrophysiology/CBV pairs to see whether inter-trial variability observed earlier (captured in the values of the CBV vectors) did correlate with the variability in behavioral or electrophysiological parameters [Figure 7B].

**Figure 7:**
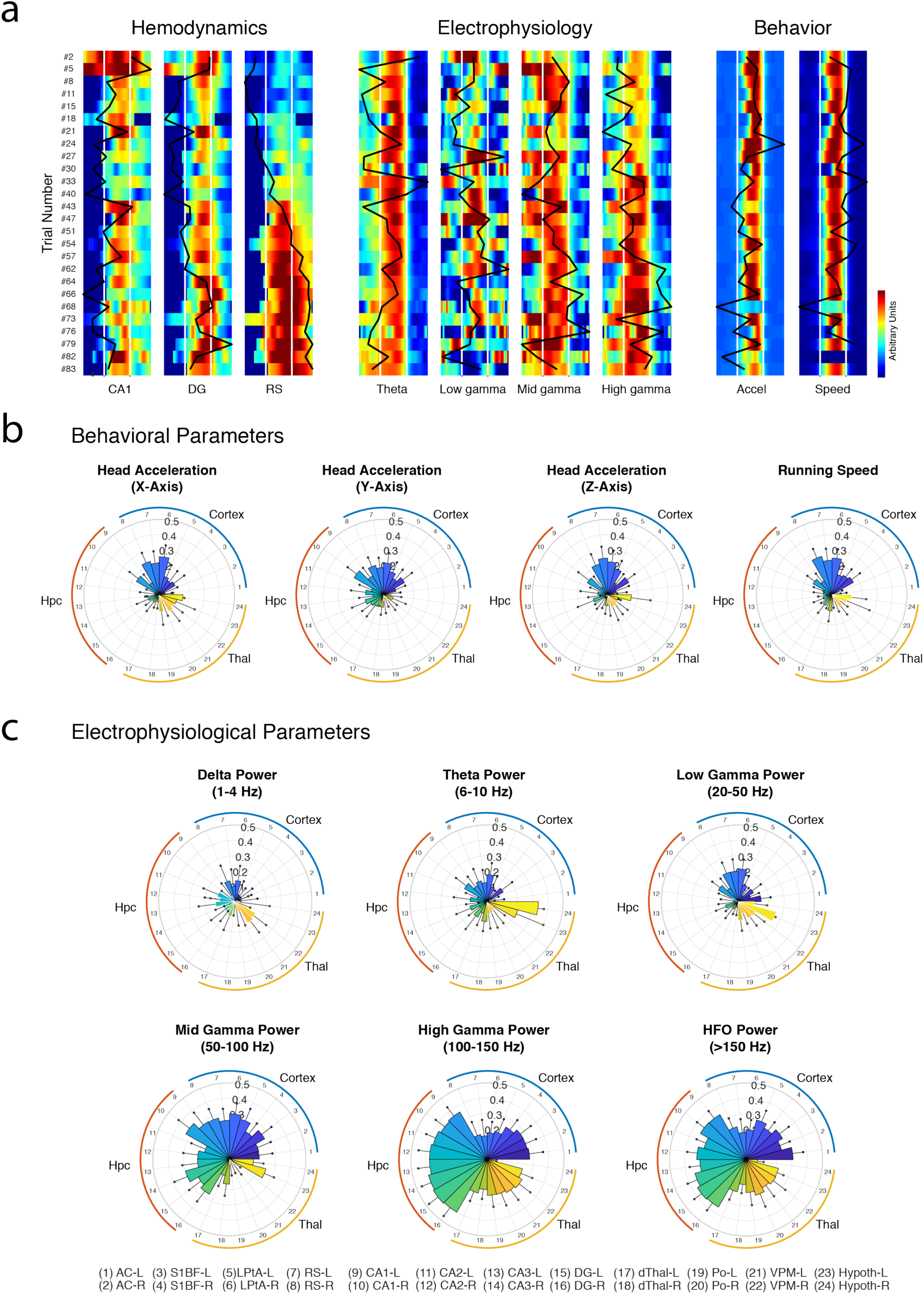
Vascular potentiation patterns are overall independent of behavior, but correlate with fast hippocampal LFP oscillations. (a) Parallel display of simultaneous CBV signals in 3 regions (CA1 region, dentate gyrus, retrosplenial) LFP power in four frequency bands (theta, low gamma, mid gamma, high gamma) and two behavioral parameters (speed, transverse head acceleration) for one recording session. Each line is a trial with corresponding run number on the left. CBV variation, LFP power, speed and acceleration power are all scaled to their minimal and maximal value over the full recording and color-coded for comparison. Trials have been re-aligned and temporally rescaled (stretched or compressed) to the same duration (4.2 seconds), white lines correspond to trial start and end. Black curves correspond to the median value for each trial. Inter-trial correlation is given by the correlation between two black curves for all LFP-CBV pairs and all behavior-CBV pairs. Note the strong linear potentiation in CA1 that correlated only with high gamma power while speed, acceleration and theta rhythm remain roughly constant. (b-c) Inter-trial correlation analysis of CBV potentiation patterns. For each recording, we analyzed whether behavioral parameters (top) or electrophysiological parameters (bottom) co-vary with CBV signal, to account for vascular potentiation patterns observed during locomotion. We thus computed a correlation coefficient between 12 behavioral/electrophysiological variables (Head acceleration (3), Running speed (1), Band-filtered LFP (6)) and 24 fUS regional hemodynamic signals. Results are displayed on polar plots: a strong correlation coefficient means that a given parameter co-varies (hence can explained) the inter-trial variability observed earlier. Black (respectively white) edges denote positive (respectively negative) correlation and bar values correspond to mean Pearson coefficient across recordings (error bars: sem). Note that neither running speed or head acceleration account for vascular plasticity in any brain regions, nor does theta (6-10 Hz) low-gamma (20-50 Hz) or mid-gamma (50-100 Hz). However, both high gamma (100-150 Hz) and HFO power (>150 Hz) correlate with vascular trial-to-trial changes, especially in the hippocampal regions, suggesting that fast oscillations may explain the strong vascular plasticity observed in the hippocampus.

We found that trial-to-trial variability was largely independent of behavioral parameters such as running speed or head acceleration in all three directions. Low frequency rhythms (theta and low gamma) did not correlate with vascular variability in any brain region. However mid gamma power correlated moderately with CBV signals especially in the cortical regions, while, interestingly, high gamma (100-150 Hz) and high-frequency oscillations (>150 Hz) correlated strongly with vascular signals in all dorsal hippocampus subfields and moderately in thalamic and cortical regions. This means that electrophysiological activity captured in middle and high-frequency oscillations might explain the potentiating patterns observed in vascular signals, while neither behavioral parameters nor theta rhythm could. Investigating recording sessions individually confirmed this assertion and we found strong correlations between high-frequency activity and vascular potentiation in the hippocampus for about half of the recordings in four different animals. Presumably, the variability in the correlation coefficients observed here can be explained by the variability in implantation sites and the fact that such high-frequency oscillations cannot easily be distinguished from spiking activity, which is hard to maintain across recordings sessions. In any case, the LFP-CBV coupling reported here is of a different nature than what was observed in previous studies^31, 34, 35^ and suggests that subtle changes in high-frequency electrophysiological activity co-vary with the strong vascular reshaping in distributed brain networks.

## Discussion

In this study, we described the vascular activations associated with natural locomotion involving a brain-wide network composed of the dorsal thalamus, parietal cortex, posterior cortex and dorsal hippocampus. Brain hemodynamics in those regions peak earlier in the dorsal thalamus and retrosplenial cortex (0.8 seconds after peaked running speed) and later in the posterior cortex and dorsal hippocampus (1.8 seconds after peaked running speed), with a consistent 300-ms delay between dentate gyrus and CA regions. We also demonstrated that brain hemodynamics were strongly reshaped between early and late runs within the same session. Some regions were rapidly depressed (cortex) while others were gradually potentiated (CA regions), while others were unaffected. These robust patterns occurred while behavioral parameters and theta rhythm were constant and could only be accounted for, in our experimental setup, by hippocampal high-frequency oscillations.

### “Online” modulation of brain activity during stereotyped behavior

Similar patterns of hemodynamic modulation, especially in the mean CBV level over slow (minute) timescale, have been reported in monkeys performing a visual task and were related to reward and to the degree of engagement in the task, high-reward trials eliciting a decrease in global CBV and low rewards showing an opposite effect^36^. This means that hemodynamic reshaping observed here could be related to longer pauses between rewards in the late part of the running task, but as animals took longer breaks towards the end of the session, CBV levels increased in the dorsal hippocampus which is the inverse effect of that seen in monkeys. Additionally, hemodynamic modulation in the cortex occurred over a rapid timescale (in the first 3 minutes) during which time between reward and running speed were remarkably constant. Importantly, hemodynamic reshaping was stronger in the dorsal hippocampus than in any other region, with a marked linear increase in CA1 regions. We do not interpret these hemodynamic changes as learning effects as these animals have been exposed to the same environment many times over days and know it quite well (overtraining). Rats are quick to learn new environments and stabilize an internal representation of space only after minutes of exploration in new mazes^37^. In our view, hemodynamic reshaping reflects ongoing plasticity processes at play during stereotyped running. They could be related to the asymmetric expansion and tuning of hippocampal place fields occurring during repeated locomotion behavior^38^. Another explanation could be that vascular networks are reorganized online during the task to reinforce relevant information location. Such reorganization of vascular networks has been shown to occur on very fast timescales to support the transfer of long-term memories from hippocampal to cortical sites in an odor-recognition task^39^. Last but not least, we observed that hemodynamic potentiation in CA1 precedes high-gamma potentiation by a few trials: one explanation could be that vascular activity directly or indirectly modulates neuronal activity in the subsequent trials, through astrocytic modulation or temperature modifications^40^.

### Complex multi-facet pattern of brain activity during running

Several studies investigated brain hemodynamics during locomotion and reported equivocal results. In head-fixed mice running on a spherical treadmill, somatosensory cortex showed a strong coupling with gamma oscillations (40-100 Hz) and firing rate, while the adjacent frontal cortex displayed prominent electrical activity with absent CBV response^21^. Arterial and venous blood showed different dynamics that contributed differently to the hemodynamic response function^41^. Studies using autoradiography radiotracers in rats performing treadmill running found changes in cerebral blood flow in dorso-lateral striatum, motor cortices and cerebellum, regions that were not imaged here^42^. Autoradiography provide a very indirect measure of brain activity and lacks sensitivity. In addition, Zhang and colleagues recently found that locomotion drives cortex-wide increases in blood oxygenation and that this effect is mediated by respiration^43^. These studies rely on the use of treadmills or similar setups which generate different patterns of electrical activity than the ones observed in natural movement, possibly because of the discrepancies between vestibular and visual sensory inputs^44^. During locomotion, we found strong activations in multiple brain regions, some of which might be associated with sensory stimulation (dorsal thalamus), spatial information (dorsal hippocampus, retrosplenial cortex), motor control (hypothalamus) or reward uptake (Po, VPM, striatum). This is consistent with a massive activation of neuronal and astrocytic networks during locomotion^20^. Diagonal planes revealed dichotomies within anatomical structures: we found different CBV profiles between ventral and dorsal hippocampus which are known to process spatial information differently^45^, between ventral and dorsal thalamic regions, both in activation onset and strength, and different timings between anterior and posterior cortices. In one individual, ventral thalamic activation occurred concurrently with frontal cortical activation during reward uptake (S1 lip region - S1 jaw) suggesting a possible thalamic relay in sensory stimulation to the face during reward uptake. These complex patterns may be associated with different components of the running task, including motor control, spatial processing, cognitive control or reward uptake. Further investigation is necessary to disentangle the functional correlates of these components, such as a comparison between reference motor-only trials and goal-directed trials, for example in a T-maze task.

### Dynamic sequence of activation in hippocampal subfields

Our recordings reveal a sequential pattern of vascular activation that originated in the dorsal thalamus and retrosplenial cortex and spread along the sub-regions of the dorsal hippocampus. The high-definition analysis of these delays revealed 250-300 milliseconds lag between dentate gyrus and CA3 peak response. These are too large to reflect the feed-forward excitation of the tri-synaptic circuit observable, for example during *in vitro* 5-Hz stimulation of the performant pathway in deafferented mice hippocampal slices. These so-called “tri-synaptic circuit waves” last between 60 and 80 milliseconds, corresponding to 5-fold faster timescale than what we observed^46^. It seems also unlikely that these patterns reflect electrical waves that propagate along the septo-temporal pole of the hippocampus as their traveling speed of 0.1-0.15 m/s is also quite high^6^. However, the fact that hippocampal subfields hemodynamic responses differ not only in their onset time but also in their temporal profile, with dentate gyrus activity starting earlier and being more sustained when CA1 activation is “steeper” and more transient, suggests that brain hemodynamics represent population activity first restricted to dentate gyrus before spreading to CA3 regions, about 300 milliseconds later. This could correspond to an excitatory stimulation of the performant path in CA3 that triggers local inhibition in the CA3 network to contain excitation via interneurons activation^47^. This interpretation also fits well with the idea that CBV signals reflect faithfully the cost of local inhibition^22^. Consistent with this idea, LTP induction via performant pathway stimulation produced detectable changes in BOLD-fMRI signals within hippocampal subfields in rats, meaning that vascular reshaping could be a proxy of local synaptic plasticity processes^48^.

### Theta, gamma and their relative role in neurovascular coupling

Previous studies that recorded concurrently cerebral hemodynamics and local field potentials have shown that fast gamma oscillation strongly correlates with subsequent vascular signals^35, 49, 50^. We recently showed that during REM sleep, phasic bouts of high gamma oscillations strongly correlate with subsequent vascular activity in almost all brain regions^31^. Using the same analysis, our present results suggest instead that during locomotion theta power – that is linearly related to speed – correlates better with CBV fluctuations than high gamma rhythms, in the dorsal thalamus, dentate gyrus and CA regions. This means that neurovascular coupling is not only region-dependent but also state-dependent. Brain hemodynamics during REM sleep were extremely synchronous in all brain regions, whereas they are more spatially decorrelated in the present study. Importantly, we found high correlations between CBV and speed or theta power in the dentate gyrus, meaning that there is a strong linear relationship between running speed, associated theta activity and subsequent vascular activation. Finally, when assessing CBV changes on slower timescales, we found that mid-gamma (50-100 Hz) correlated with CBV fluctuations in cortical regions whereas high-gamma or high-frequency oscillations (100-150 Hz) did so in the dorsal hippocampus and thalamus. Concurrently, low-gamma activity elicited weak to no vascular response in most brain regions. This strongly suggests the existence of distinct vascular network areas preferentially associated with different frequency bands, which respond to activity within these bands over different timescales.

### Other parameters that could drive hemodynamic reshaping

Several important parameters could modulate brain hemodynamics during locomotion including breathing, heart rate, respiration or temperature. Controlling all these parameters independently remains to be done in the future, but we can already rule out or at least lower the probable influence of these parameters based on the spatial profile of hemodynamic reshaping. A major homeostatic effect arising from increased heart rate, respiration or neuromodulation is likely to affect all brain structures similarly: we would probably not observe a decrease in the cortex, concurrent with an increase in the CA1 region and a stable response in the dorsal thalamus if heart rate was the main driver of hemodynamic reshaping. Also, these parameters are directly related to running speed which remains constant across trials. Brain temperature however is known to affect brain activity significantly^51^ and it is likely to increase with intrinsic heat generated by muscular effort over the accumulation of trials, together with the direct ultrasound-insonification. But because we use a sequence that imposes 40-second refractory period (resulting in a 20-25% efficacy imaging time) and the fact that our animals underwent large craniotomies should result in a substantial dissipation of heat in brain tissue^52^. Also, neighboring structures like dorsal thalamus and ventral thalamus, or dentate gyrus and CA1 region show strong discrepancies in their modulation pattern, thus if we cannot completely exclude an effect of temperature, it probably superimposes on other underlying causes of vascular reshaping.

One of the conclusions of our study is that inter-trial averaging must be performed with caution (for example over a small number of trials) even when behavioral or coarse electrophysiological parameters such as theta rhythms suggest that two given trials are similar. Second, it shows that the neurovascular interactions are complex and couple different regions with different frequency bands over multiple timescale: though hippocampal theta rhythm reports vascular activity in the CA1 region at the second timescale pretty faithfully, it fails to account for the slower modifications of CA1 hemodynamics across trials, which are massive. Probing the activity of brain hemodynamics at large-scale and comparing to local neuronal activity monitored with electrodes is a work in progress that will considerably help us understand the physiological basis of complex behavior.

## Online Methods

### Animal Surgery

All animals received humane care in compliance with the European Communities Council Directive of 2010 (2010/63/EU), and the institutional and regional committees for animal care approved the study. Adult Sprague Dawley rats aged 10-12 weeks underwent surgical craniotomy and implant of an ultrasound-clear prosthesis. Anesthesia was induced with 2% isoflurane and maintained with ketamine/xylazine (80/10 mg/kg), while body temperature was maintained at 36.5°C with a heating blanket (Bioseb, France). A sagittal skin incision was performed across the posterior part of the head to expose the skull. We excised the parietal and frontal flaps by drilling and gently moving the bone away from the dura mater. The opening exposed the brain between the olfactory bulb and the cerebellum, from Bregma +6.0 to Bregma −8.0 mm, with a maximal width of 14 mm. Electrodes were implanted stereotaxically and anchored on the edge of the flap. A prosthetic skull was sealed in place with acrylic resin (GC Unifast TRAD), and the residual space was filled with saline. We chose a prosthesis approach that offers a larger field of view and prolonged imaging condition over 4-6 weeks compared to the thinned bone approach. The prosthetic skull is composed of polymethylpentene (Goodfellow, Huntington UK, goodfellow.com), a standard biopolymer used for implants. This material has tissue-like acoustic impedance that allows undistorted propagation of ultrasound waves at the acoustic gel-prosthesis and prosthesis-saline interfaces. The prosthesis was cut out of a film of 250 µm thickness and permanently sealed to the skull. Particular care was taken not to tear the dura to prevent cerebral damage. The surgical procedure, including electrode implantation, typically took 4-6 hours. Animals recovered quickly and were used for data acquisition after a conservative one-week resting period.

### Electrode design and implantation

Electrodes are based on linear polytrodes made of bundles of insulated tungsten wires. The difference with a standard design is a 90°-angle elbow that is formed prior to insertion in the brain^28^. This shape enabled anchoring of the electrodes on the skull anterior or posterior to the flap. Electrodes were implanted with stereotaxic positioning micromotion and anchored one after another. The prosthesis was then applied to seal the skull. Two epidural screws placed above the cerebellum were used as a reference and ground. Intra-hippocampal handmade electrode bundles were composed of 25 to 50 µm diameter insulated tungsten wire soldered to miniature connectors. Four to six conductive ends were spaced 1 mm apart and glued to form 3-mm-long, 100-150-µm-diameter bundles. The bundles were lowered in the dorsal hippocampi at stereotaxic coordinates AP = −4.0 mm, ML = +/- 2.5 mm and DV = −1.5 mm to −4.5 mm relative to the Bregma.

### LFP acquisition

LFP signals were collected from video-EEG device for offline processing. Intracranial electrode signals were fed through a high input impedance, DC-cut at 1 Hz, gain of 1000, 16-channel amplifier and digitized at 20 kHz (Xcell, Dipsi, Cancale, France), together with a synchronization signal from the ultrasound scanner. Custom-made software based on LabVIEW (National Instruments, Austin, TX, USA) simultaneously acquired video from a camera pointed at the recording stage. A regular amplifier was used, and no additional electronic circuit for artifact suppression was necessary. A large bandwidth amplifier was used, which can record local field potentials in all physiological bands (LFP, 0.1-2 kHz). The spatial resolution of LFPs ranges from 250 µm to a few mm radius.

### Ultrasound Acquisition

Vascular images were obtained via the ultrafast compound Doppler imaging technique^53^. The probe was driven by a fully programmable GPU-based ultrafast ultrasound scanner (Aixplorer, Supersonic Imagine, Aix-en-Provence, France) relying on 24-Gb RAM memory. We acquired 6000 ultrasound images at 500 Hz frame rate for 12 seconds, repeating every 52-60 s (refractory period of 40 seconds during which no data can be acquired). Each frame is a compound plane-wave frame, that is, a coherent summation of beamformed complex in phase/quadrature images obtained from the insonification of the medium with a set of successive plane waves with specific tilting angles^54^. This compound plane wave imaging technique enabled the re-creation of a dynamic transmit focusing at all depths *a posteriori* in the entire field of view with few ultrasound emissions. Given the tradeoff between frame rate, resolution and imaging speed, a plane-wave compounding using five 5°-apart angles of insonification (from −10° to +10°) was chosen. As a result, the pulse repetition frequency of the plane wave transmissions was equal to 500 Hz. To discriminate blood signals from tissue clutter, the ultrafast compound Doppler frame stack was filtered, removing the N = 60 first components of the singular value decomposition, which optimally exploited the spatiotemporal dynamics of the full Doppler film for clutter rejection, largely outperforming conventional clutter rejection filters used in Doppler ultrasound^55^.

### Recording sessions

Eleven male Sprague Dawley were trained daily to run back and forth on a 2.2-meter linear track for a water reward given on both ends through solenoid valves. The rats were placed under a controlled water restriction protocol (weight maintained between 85 and 90% of the normal weight) and trained for one week before surgery. The track (225 x 20 cm) had 5-cm-high lateral walls and was placed 50 cm above the ground. Each time the animal crossed the middle of the track, one drop of water was delivered in alternate water tubes by opening an electronically controlled pair of solenoid valves. Daily training lasted 30 min. Rats then underwent a surgery combining implantation of handmade linear LFP probes in the dorsal hippocampus with permanent skull replacement by a polymer prosthesis that allows undistorted ultrasound passage. At the start of each recording session, to attach the ultrasound probe and connect the EEG, the rats underwent brief anesthesia for 20-25 min with 2% isoflurane. Acoustic gel was applied on the skull prosthesis, and the probe was inserted into the probe holder. The gel did not dry out even for extended recordings of up to 6-8 hours. The animals were allowed to recover for 40 min before starting the recording session. A typical session included a 30-40 min running period. In total, we recorded from 11 animals over the coronal and diagonal planes for a total of 42 running sessions. Though overall distance traveled was lower in mfUS-EEG animals than in controls, our protocol did not impede instantaneous animal movement (no significant difference in peak speed). This can be explained by the higher load onto the animal’s head, resulting in longer inter-trial interval, probably due to longer rest periods^28^.

### LFP/VIDEO analysis

All analysis was performed in MATLAB. For each recording, the position of each recording site on the probe tract (four recording sites per probe) was identified by measuring its impedance while lowering the bundle in saline solution (Na-Cl 0.9%) before implantation. Hippocampal theta and gamma rhythms were confirmed by phase inversion across recording sites in successive hippocampal layers, time-frequency decomposition and phase-amplitude cross-frequency coupling^18^. Though we cannot completely exclude potential contamination of high-frequency LFP recordings by unit activity, we observed that fast gamma oscillations displayed typical phase-amplitude coupling patterns for all recordings robustly across animals. The size of our electrode diameters (25 to 50 microns) and the stability of our recordings throughout sessions decrease the probability that fast gamma events arose from correlated spiking activity. We then selected the putative CA1 and dentate gyrus recording sites based on theta-gamma coupling patterns and computed a differential signal from these two traces for each animal. EEG was first filtered in the LFP range (LFP, 0.1-2 kHz) and band-pass filtered in typical frequency bands including delta (1-4 Hz), theta (6-10 Hz), low-gamma (20-50 Hz), mid-gamma (50-100 Hz), high-gamma (100-150 Hz) and ripple (150-250 Hz). This division has been thoroughly described and proven to be functionally relevant for hippocampal electrographic recordings^33^. The power of LFP oscillations was computed as the square of the raw signal integrated over a sliding Gaussian kernel of a characteristic width of 500 milliseconds to extract its envelope. To select the putative CA1 recording site, we computed the instantaneous theta phase (taking 0° as a theta trough) using a Hilbert transform and derived theta-phase time-frequency spectrograms to exhibit phase-amplitude theta-gamma coupling patterns. CA1 recording sites were identified when peak gamma power occurs at the peak of the theta phase, corresponding to the CA1 *stratum lacunosum-moleculare*^18^. Videos were processed frame by frame, first by drawing a fixed rectangle to isolate all pixels within the track. We then performed thresholding on the histogram of the pixels inside the track to isolate bright pixels of the rat’s body and compute their barycenter. Animal’s position is then computed frame by frame and smoothed with a gaussian kernel (half-width 500 ms). Speed vectors are then extracted in two dimensions and re-projected into the actual track coordinate to extract relevant body speed.

### Baseline evaluation & spatial averaging

fUS data have been shown to be proportional to local CBV. Because it is not possible to derive absolute CBV levels, we normalized ultrasound data. We performed voxel-wise normalization from a baseline period: before the animal started the session, we systematically recorded three 12-second lapses during quiet wakefulness while the animal’s cage was placed nearby one end of the track. We extracted the distribution for each voxel during this baseline period and computed a mean value, leading to one value for each voxel of the image. To derive a signal similar to ΔF/F, we subtracted the mean and divided by the mean for each voxel in the film containing our power Doppler images. This allowed normalization and rescaling of ultrasound data, yielding to an expression in terms of the percent of variation relative to baseline. Each voxel was normalized independently before performing spatial averaging.

### Atlas registration

Coronal recordings were registered to two-dimensional sections from the Paxinos atlas^56^ using anatomical landmarks, such as cortex edges, hippocampus outer shape and large vessels below brain surface as a reference. We performed manual scaling and rotation along each of the 3 dimensions to recover the most probable registration. Once performed, regions of interest were extracted using binary masks. To register diagonal planes that had no direct coronal sections, we segmented each two-dimensional recording plane into anatomical regions based on a 3D MRI-based whole-brain atlas, which provided labeling for 52 brain regions^57^. To derive the functional regions in our ultrasound image, we designed a customized registration algorithm^31^, which projected our two-dimensional ultrasound plane onto a three-dimensional volumetric dataset. In short, we manually pinpointed landmarks on the ultrasound image including the outer cortex edges, inner cortex edges, midline plane and dentate gyri edges, which were prominent due to hippocampal longitudinal arteries wrapping them. We defined 9 parameters, including 3 offset values, 3 scaling values and 3 rotations (13 parameters for multiplane registration), to identify a given plane unambiguously in the 3D Waxholm space. We performed optimization using the simplex algorithm to minimize a global error based on the position of our landmarks and the closest corresponding border in the Waxholm space. Provided the algorithm did not start too far from the actual position, it converged quickly and provided robust registration for any ultrasound plane, including diagonal planes. This process allowed us to derive vascular activity in 20 regions that were observed in two imaging planes intersecting the recording electrode tracts (one coronal view at coordinates Bregma=- 4.0 mm and one diagonal plane 45° relative to the sagittal plane to include the ventral and dorsal hippocampus).

### LFP-CBV correlation analysis

To assess the association between LFP events and CBV variables, we searched for correlations between each possible combination of one LFP band-pass filtered signal and a given regional CBV variable. As neurovascular processes are not instantaneous, we considered possible delays between electrographic and vascular signals. We thus directly computed lagged cross-correlations between two signals for any LFP-CBV pair and any lag in a given time window. We also performed this over regional variables to perform statistical comparison across recordings.

### Statistics

All statistics are given as +/- standard error of the mean unless stated otherwise. Statistics in Fig 3 are computed on n=11 animals over 42 recording sessions. Bar diagrams shown in Fig 3 are computed by averaging the mean values of 22 recordings on 11 animals for the coronal planes and 20 recordings in 7 animals for the diagonal planes. Statistics are computed using a two-tailed Mann-Whitney test. The significance of Pearson correlation coefficients shown in Fig 3 and Fig 7 are assessed by computing the t-value (using 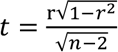) and reporting it in Student’s table with n-2 degrees of freedom.

## Acknowledgements

The authors would like to thank S. Hubatz for help with the track running protocol, T. Watson for technical help with current-lesioning protocol. A.B. received funding from the Ecole Doctorale Frontières du Vivant, Programme Bettencourt. The research leading to these results has received funding from the European Research Council under the European Union’s Seventh Framework Programme (*FP7/2007-2013)* / ERC *grant agreement* n° 339244-FUSIMAGINE. This work was also partly supported by the Fondation pour la Recherche sur le Cerveau (FRC) (Program Rotary – Espoir en tête) and by the AXA Research Fund under the chair *New hopes in medical imaging with ultrasound*.

## Author contributions

A.B. and I.C. designed the experiment. A.B. designed the electrodes, performed the surgeries, training and recording sessions. E.T. and T.D. programmed the ultrasound sequences and burst recording mode. C.D. designed the clutter-rejection algorithm. I.C. programmed the acquisition software and atlas-registration algorithm. A.B. analyzed the behavioral and electrographic data. A.B. and M.T. analyzed the ultrasound data and discussed multimodal analysis. All authors wrote the paper.

## List of Figures

### Supplementary Figures

**Figure S1:** Large-scale hemodynamic responses of locomotion in freely-running rats (diagonal plane)

**Figure S2:** Comparison of hemodynamic responses with respect to running direction

**Figure S3:** Details of inter-trial variability for all recordings

**Figure S4:** Details of vascular potentiation for all recordings

**Figure S5:** Comparative analysis of seed-based correlations (Coronal plane recordings)

**Figure S6:** Comparative analysis of seed-based correlations (Diagonal plane recordings)

### Supplementary Movies

**Supplementary Movie 1-2:** Sequential activation of the tri-synaptic circuit at the second timescale over coronal and diagonal recording planes

**Supplementary Movie 3-4:** Task-related hippocampal potentiation and cortical depression throughout a single recording during locomotion

## References

1. Berger, H. Über das elektrenkephalogramm des menschen. European Archives of Psychiatry and Clinical Neuroscience 87, 527–570 (1929).

2. Buzsáki, G. Rhythms of the brain. (Oxford Univ. Press, 2006).

3. Fries, P. A mechanism for cognitive dynamics: neuronal communication through neuronal coherence. Trends in Cognitive Sciences 9, 474–480 (2005).

4. Fell, J. & Axmacher, N. The role of phase synchronization in memory processes. Nature Reviews Neuroscience 12, 105–118 (2011).

5. Schnitzler, A. & Gross, J. Normal and pathological oscillatory communication in the brain. Nature Reviews Neuroscience 6, 285–296 (2005).

6. Lubenov, E. V. & Siapas, A. G. Hippocampal theta oscillations are travelling waves. Nature 459, 534–539 (2009).

7. Patel, J., Fujisawa, S., Berényi, A., Royer, S. & Buzsáki, G. Traveling Theta Waves along the Entire Septotemporal Axis of the Hippocampus. Neuron 75, 410–417 (2012).

8. Massimini, M. The Sleep Slow Oscillation as a Traveling Wave. Journal of Neuroscience 24, 6862–6870 (2004).

9. Muller, L., Chavane, F., Reynolds, J. & Sejnowski, T. J. Cortical travelling waves: mechanisms and computational principles. Nature Reviews Neuroscience 19, 255–268 (2018).

10. Buzsáki, G. Theta oscillations in the hippocampus. Neuron 33, 325–340 (2002).

11. Colgin, L. L. Rhythms of the hippocampal network. Nature Reviews Neuroscience 17, 239–249 (2016).

12. Bland, B. H. & Oddie, S. D. Theta band oscillation and synchrony in the hippocampal formation and associated structures: the case for its role in sensorimotor integration. Behavioural brain research 127, 119–136 (2001).

13. Jezek, K., Henriksen, E. J., Treves, A., Moser, E. I. & Moser, M.-B. Theta-paced flickering between place-cell maps in the hippocampus. Nature 478, 246–249 (2011).

14. Benchenane, K. et al. Coherent Theta Oscillations and Reorganization of Spike Timing in the Hippocampal-Prefrontal Network upon Learning. Neuron 66, 921–936 (2010).

15. Boyce, R., Glasgow, S. D., Williams, S. & Adamantidis, A. Causal evidence for the role of REM sleep theta rhythm in contextual memory consolidation. Science 352, 812–816 (2016).

16. Drieu, C., Todorova, R. & Zugaro, M. Nested sequences of hippocampal assemblies during behavior support subsequent sleep replay. 6 (2018).

17. Jensen, O. & Colgin, L. L. Cross-frequency coupling between neuronal oscillations. Trends in Cognitive Sciences 11, 267–269 (2007).

18. Schomburg, E. W. et al. Theta Phase Segregation of Input-Specific Gamma Patterns in Entorhinal-Hippocampal Networks. Neuron 84, 470–485 (2014).

19. Colgin, L. L. et al. Frequency of gamma oscillations routes flow of information in the hippocampus. Nature 462, 353–357 (2009).

20. Dombeck, D. A., Khabbaz, A. N., Collman, F., Adelman, T. L. & Tank, D. W. Imaging Large-Scale Neural Activity with Cellular Resolution in Awake, Mobile Mice. Neuron 56, 43–57 (2007).

21. Huo, B.-X., Smith, J. B. & Drew, P. J. Neurovascular Coupling and Decoupling in the Cortex during Voluntary Locomotion. Journal of Neuroscience 34, 10975–10981 (2014).

22. Buzsáki, G., Kaila, K. & Raichle, M. Inhibition and Brain Work. Neuron 56, 771–783 (2007).

23. Attwell, D. et al. Glial and neuronal control of brain blood flow. Nature 468, 232–243 (2010).

24. Kleinfeld, D. et al. A Guide to Delineate the Logic of Neurovascular Signaling in the Brain. Frontiers in Neuroenergetics 3, (2011).

25. Devonshire, I. M. et al. Neurovascular coupling is brain region-dependent. NeuroImage 59, 1997–2006 (2012).

26. Lin, A.-L., Fox, P. T., Hardies, J., Duong, T. Q. & Gao, J.-H. Nonlinear coupling between cerebral blood flow, oxygen consumption, and ATP production in human visual cortex. PNAS 107, 8446–8451 (2010).

27. Uhlirova, H. et al. Cell type specificity of neurovascular coupling in cerebral cortex. eLife 5, e14315 (2016).

28. Sieu, L.-A. et al. EEG and functional ultrasound imaging in mobile rats. Nature Methods 12, 831–834 (2015).

29. Sauvage, J. et al. A large aperture row column addressed probe for *in vivo* 4D ultrafast doppler ultrasound imaging. Physics in Medicine & Biology 63, 215012 (2018).

30. Rabut, C. et al. 4D functional ultrasound imaging of whole-brain activity in rodents. Nature Methods 16, 994–997 (2019).

31. Bergel, A., Deffieux, T., Demené, C., Tanter, M. & Cohen, I. Local hippocampal fast gamma rhythms precede brain-wide hyperemic patterns during spontaneous rodent REM sleep. Nature Communications 9, (2018).

32. Macé, E. et al. Functional ultrasound imaging of the brain. Nature Methods 8, 662–664 (2011).

33. Belluscio, M. A., Mizuseki, K., Schmidt, R., Kempter, R. & Buzsaki, G. Cross-Frequency Phase-Phase Coupling between Theta and Gamma Oscillations in the Hippocampus. Journal of Neuroscience 32, 423–435 (2012).

34. Bouchard, M. B., Chen, B. R., Burgess, S. A. & Hillman, E. M. C. Ultra-fast multispectral optical imaging of cortical oxygenation, blood flow, and intracellular calcium dynamics. Opt Express 17, 15670–15678 (2009).

35. Mateo, C., Knutsen, P. M., Tsai, P. S., Shih, A. Y. & Kleinfeld, D. Entrainment of Arteriole Vasomotor Fluctuations by Neural Activity Is a Basis of Blood-Oxygenation-Level-Dependent “Resting-State” Connectivity. Neuron 96, 936–948.e3 (2017).

36. Cardoso, M. M. B., Lima, B., Sirotin, Y. B. & Das, A. Task-related hemodynamic responses are modulated by reward and task engagement. PLOS Biology 17, e3000080 (2019).

37. O’Keefe, J. & Nadel, L. The hippocampus as a cognitive map. (Clarendon Press, 1978).

38. Mehta, M. R., Barnes, C. A. & McNaughton, B. L. Experience-dependent, asymmetric expansion of hippocampal place fields. PNAS 94, 8918–8921 (1997).

39. Frankland, P. W. & Bontempi, B. The organization of recent and remote memories. Nature Reviews Neuroscience 6, 119–130 (2005).

40. Moore, C. I. & Cao, R. The Hemo-Neural Hypothesis: On The Role of Blood Flow in Information Processing. Journal of Neurophysiology 99, 2035–2047 (2008).

41. Huo, B.-X., Gao, Y.-R. & Drew, P. J. Quantitative separation of arterial and venous cerebral blood volume increases during voluntary locomotion. NeuroImage 105, 369–379 (2015).

42. Holschneider, D. P. & Maarek, J.-M. I. Mapping brain function in freely moving subjects. Neuroscience & Biobehavioral Reviews 28, 449–461 (2004).

43. Zhang, Q. et al. Cerebral oxygenation during locomotion is modulated by respiration. Nature Communications 10, (2019).

44. Li, J.-Y., Kuo, T. B. J., Yen, J.-C., Tsai, S.-C. & Yang, C. C. H. Voluntary and involuntary running in the rat show different patterns of theta rhythm, physical activity, and heart rate. Journal of Neurophysiology 111, 2061–2070 (2014).

45. Strange, B. A., Witter, M. P., Lein, E. S. & Moser, E. I. Functional organization of the hippocampal longitudinal axis. Nature Reviews Neuroscience 15, 655–669 (2014).

46. Stepan, J., Dine, J. & Eder, M. Functional optical probing of the hippocampal trisynaptic circuit in vitro: network dynamics, filter properties, and polysynaptic induction of CA1 LTP. Frontiers in Neuroscience 9, (2015).

47. Lawrence, J. J. & McBain, C. J. Interneuron Diversity series: Containing the detonation – feedforward inhibition in the CA3 hippocampus. Trends in Neurosciences 26, 631–640 (2003).

48. Abe, Y., Tsurugizawa, T., Le Bihan, D. & Ciobanu, L. Spatial contribution of hippocampal BOLD activation in high-resolution fMRI. Scientific Reports 9, (2019).

49. Logothetis, N. K., Pauls, J., Augath, M., Trinath, T. & Oeltermann, A. Neurophysiological investigation of the basis of the fMRI signal. Nature 412, 150–157 (2001).

50. Niessing, J. Hemodynamic Signals Correlate Tightly with Synchronized Gamma Oscillations. Science 309, 948–951 (2005).

51. Moser, E. I. & Mathiesen, lacob. Relationship between neuronal activity and brain temperature in rats: NeuroReport 7, 1876 (1996).

52. Roche, M. et al. In vivo imaging with a water immersion objective affects brain temperature, blood flow and oxygenation. eLife 8, (2019).

## Methods-only references

53. Tanter, M. & Fink, M. Ultrafast imaging in biomedical ultrasound. *IEEE Transactions on Ultrasonics*, Ferroelectrics, and Frequency Control 61, 102–119 (2014).

54. Montaldo, G., Tanter, M., Bercoff, J., Benech, N. & Fink, M. Coherent plane-wave compounding for very high frame rate ultrasonography and transient elastography. *IEEE Transactions on Ultrasonics*, Ferroelectrics and Frequency Control 56, 489–506 (2009).

55. Demene, C. et al. Spatiotemporal Clutter Filtering of Ultrafast Ultrasound Data Highly Increases Doppler and fUltrasound Sensitivity. IEEE Transactions on Medical Imaging 34, 2271–2285 (2015).

56. Paxinos, G. & Watson, C. The Rat Brain in Stereotaxic Coordinates. (Academic Press, 1982).

57. Papp, E. A., Leergaard, T. B., Calabrese, E., Johnson, G. A. & Bjaalie, J. G. Waxholm Space atlas of the Sprague Dawley rat brain. NeuroImage 97, 374–386 (2014).

58. LaMendola, N. P. & Bever, T. G. Peripheral and cerebral asymmetries in the rat. Science 278, 483–486 (1997).

