## Supplementary Information for "“Online” modulation of brain hemodynamics despite stereotyped running"

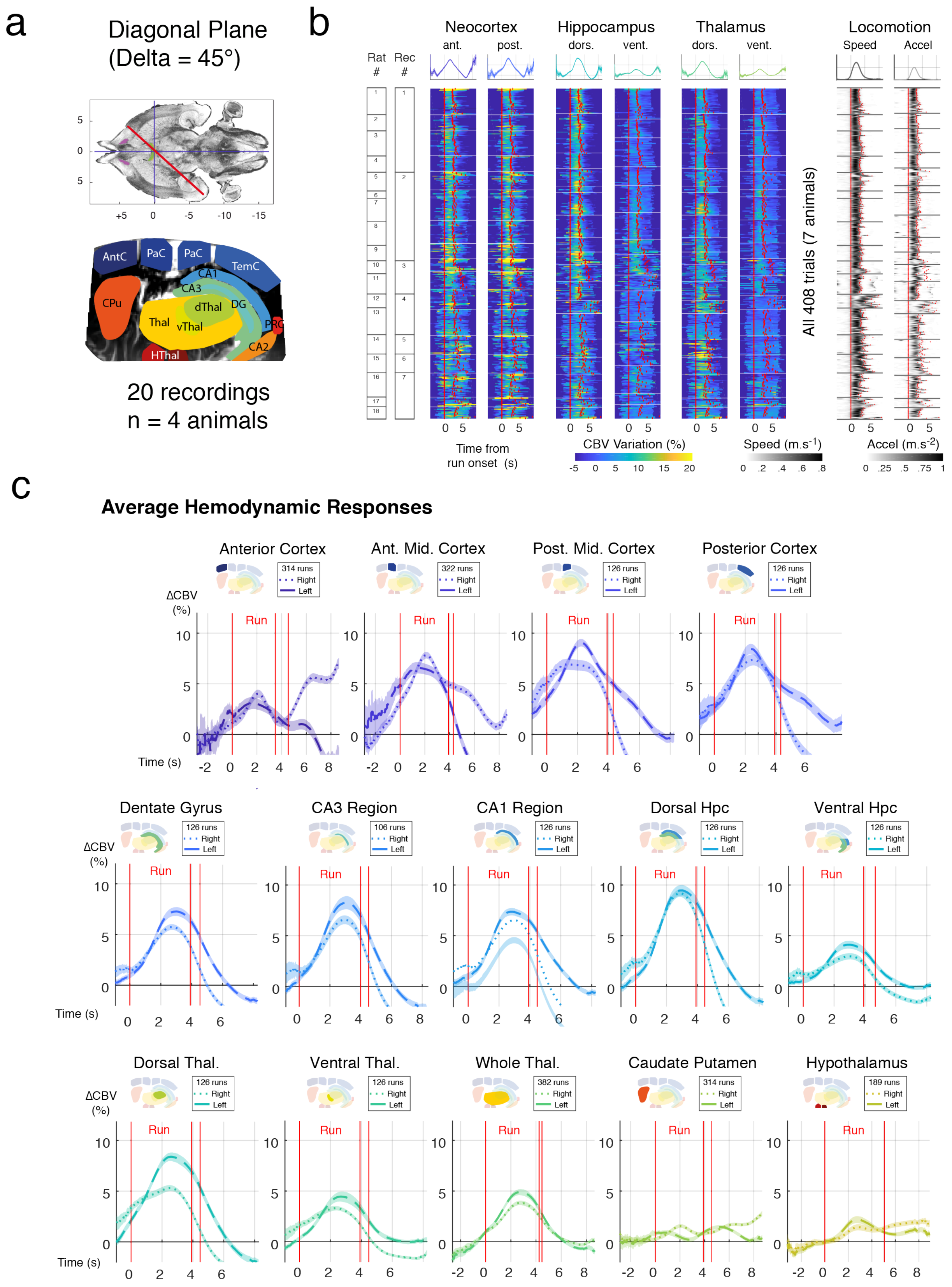


**Figure S1: Large-scale hemodynamic responses of locomotion in freely-running rats (diagonal plane)**

(a) Location of a typical diagonal (Delta = 45°) recording plane and associated brain structures monitored during mfUS-EEG recordings. Top: Plane position (red) on a transverse view of rat brain atlas. Bottom Power Doppler image with superimposed atlas registration, performed by positioning salient landmarks onto Power Doppler image and registering a 3D volumetric segmented MRI atlas. AntC: anterior cortex, PaC: parietal cortex, PostC: posterior cortex, DG: dentate gyrus, CA1-CA2-CA3 region, dThal: dorsal Thalamus, vThal: ventral Thalamus, CPu: caudate Putamen, Hthal: Hypothalamus.

(b) Bulk representation of single-trial hemodynamic responses to locomotion in six major brain regions (anterior/posterior Neocortex, ventral/dorsal Hippocampus, ventral/dorsal Thalamus). A total of 408 trials (19 recordings, 7 animals) was acquired across many days. For each run, the onset of movement is used as a temporal reference (zero-timing) and all trials are aligned to run onset (see Methods). Run end however is different for all trials. We can then compute average hemodynamic responses (top) and average run duration (t = 4.26s +/- .3s). The same approach is performed for locomotion parameters: running speed (left) and accelerometer (right). Hemodynamic responses display strong inter-trial variability in the cortical and hippocampal regions, with a strong dissociation between dorsal versus ventral hippocampus and dorsal versus ventral thalamic regions.

(c) Average hemodynamics responses over multiple sub-regions. Using the approach in b, we computed average responses to locomotion in 26 regions. Overall, strong bilateral activations are found in the cortex and all dorsal hippocampus sub-regions (stronger and earlier in the dentate gyrus), together with dorsal thalamus. We display hemodynamic responses for large regions as a reference on the right side. For each region the number of runs can differ slightly as all 26 regions were not always visible on each recording.

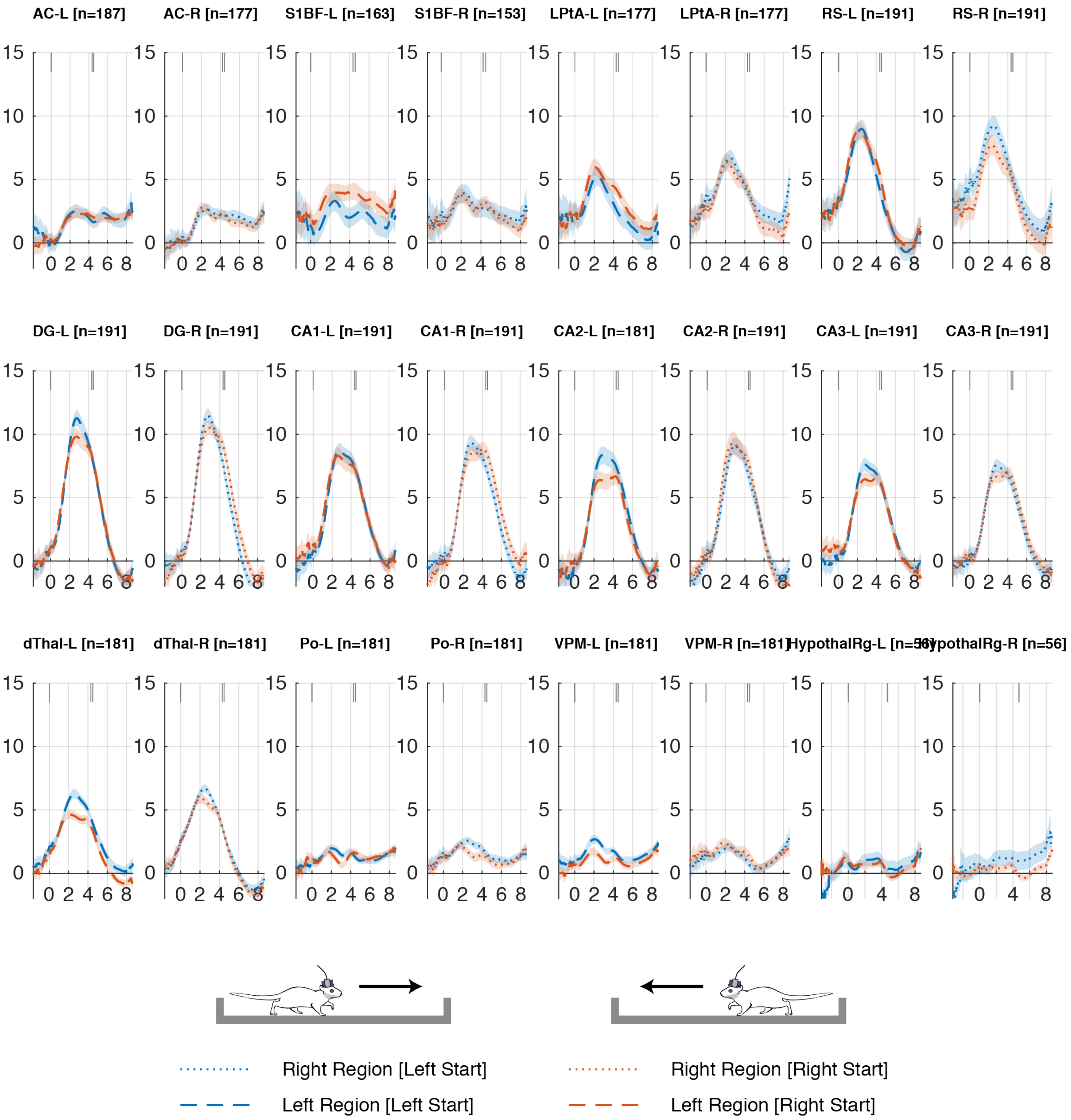


**Figure S2: Comparison of hemodynamic responses with respect to running direction**

Average hemodynamic responses for all coronal recordings segregated according each running direction (n= 7 animals, 19 recordings, 191 left runs, 191 right runs for a total of 388 runs). For each region, the average hemodynamic response for left runs is displayed in blue while the hemodynamic response to right runs is displayed in orange. Brain hemodynamics display very little dependence on running direction both in terms of delay, duration and amplitude. Non-statistically significant differences are visible in the peak amplitude of left dentate gyrus, left CA2 region, left CA3 region and left dorsal thalamus but they were largely influenced by one recording where responses were unilateral. This did not appear on other recordings. More prominent differences are observed within cortical regions (right retrosplenial, left somatosensory, left parietal associative) which might be due to asymmetries during running and reward uptake as some animals showed a preferred rotation side and ran closer to one wall compared to the other[58].

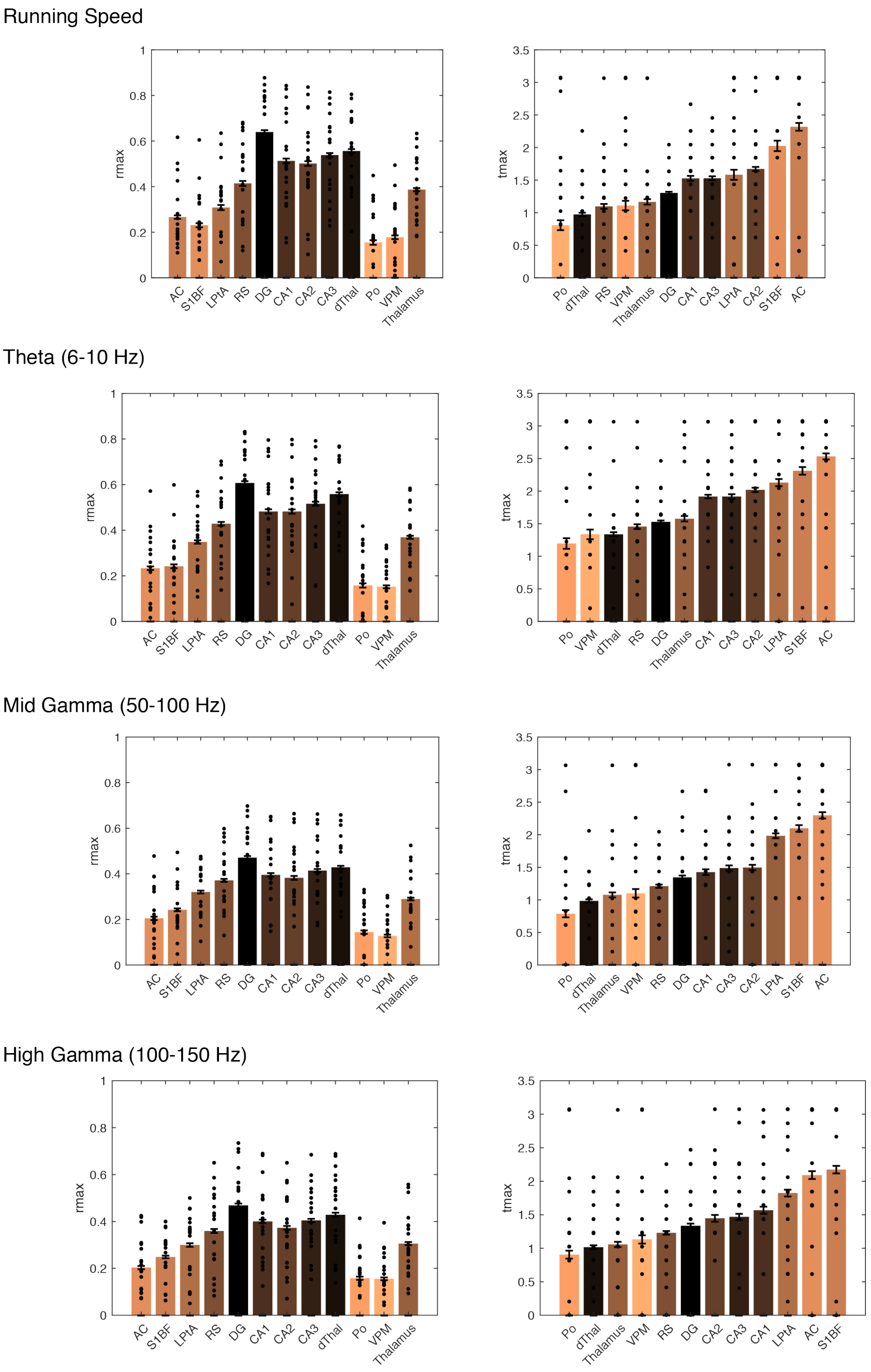


**Figure S3: Comparative analysis of seed-based correlations (Coronal recordings)**

We repeated the correlation analysis presented in Figure 3 over all coronal recordings, but changed the reference variable to compute cross-correlations. In the main text, we computed speed-CBV cross-correlations and extracted $R_{max}$ and $T_{max}$ for all CBV regional response (top), which provides a measure of the proportion of variance in CBV fluctuations that is explained by speed. Here, we also computed LFP-CBV cross-correlation for 3 different LFP envelope signals (from top to bottom): theta (6-10 Hz), mid gamma (50-100 Hz) and high gamma (100-150 Hz) and displayed the average $R_{max}$ (left) and $T_{max}$ (right) values for all recordings in all 12 regions. In the right display, regions are ranked in ascending order relative to $T_{max}$. We found that speed and theta best correlate with CBV signals in all regions, compared to mid and high-gamma power. Interestingly, the sequence of activation revealed by $T_{max}$ is remarkably well-conserved for all 4 analyses, meaning that it reveals the intrinsic dynamics of CBV regional activations independent of the reference variable used.

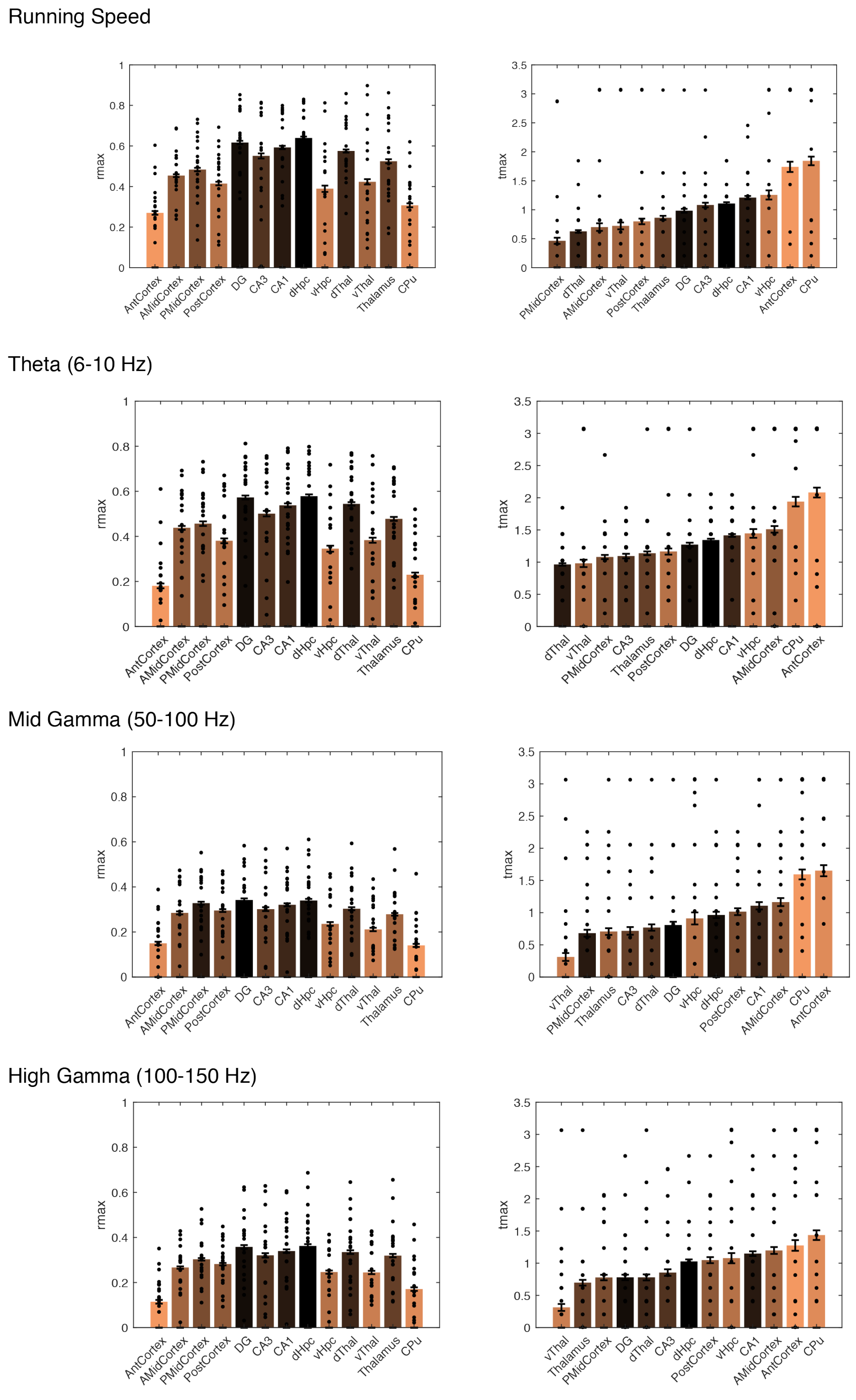


**Figure S4: Comparative analysis of seed-based correlations (Diagonal plane recordings)**

We repeated the correlation analysis presented in Figure 3 over all diagonal recordings, but changed the reference variable to compute cross-correlations. In the main text, we computed speed-CBV cross-correlations and extracted $R_{max}$ and $T_{max}$ for all CBV regional responses (top), which provides a measure of the proportion of variance in CBV fluctuations that is explained by speed. Here, we also computed LFP-CBV cross-correlation for 3 different LFP envelope signals (from top to bottom): theta (6-10 Hz), mid gamma (50-100 Hz) and high gamma (100-150 Hz) and displayed the average $R_{max}$ (left) and $T_{max}$ (right) values for all recordings in all 12 regions. In the right display, regions are ranked in ascending order relative to $T_{max}$. We found that speed and theta best correlate with CBV signals in all regions, compared to mid and high-gamma power. Interestingly, the sequence of activation revealed by $T_{max}$ is remarkably well-conserved for all 4 analyses, meaning that it reveals the intrinsic dynamics of CBV regional activations independent of the reference variable used.

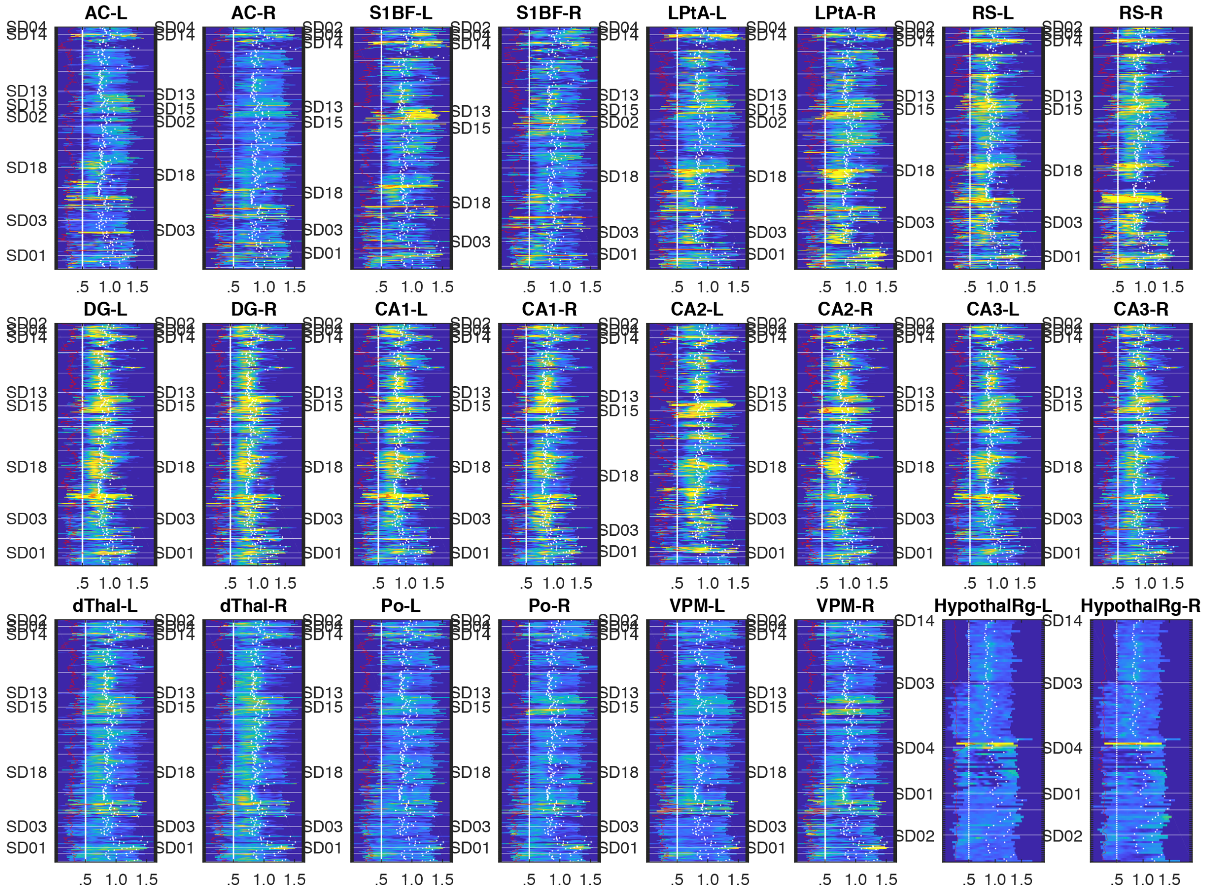


**Figure S5: Brain-wide hemodynamic modulation is directly observable in individual responses to single runs.**

Bulk representation of single-trial hemodynamic responses to locomotion in all brain regions recorded on the coronal plane (Anteroposterior axis: Bregma = - 4.0 mm) (RS: retrosplenial cortex, LPtA: lateral parietal associative cortex, S1BF: S1 barrel field, AC: auditory cortex, DG: dentate gyrus, CA1-CA2-CA3 region, dThal: dorsal Thalamus, Po: Posterior Thalamic Nucleus, VPM: Ventroposterior Thalamic Nucleus, Thal: Thalamus, Hthal: Hypothalamus). A total of 384 trials (19 recordings, 7 animals) was acquired across many days. For each run, the onset of movement is used as a temporal reference (zero-timing) and all trials are aligned to run onset (see Methods). Strongest activations are found in all subfields of the dorsal hippocampus and retrosplenial and lateral parietal cortices. Recording sessions are separated by horizontal grey lines and grouped by individuals. Animal identifier is given on the left. Note the strong inter-trial variability within the same recording session in hippocampal and cortical regions, with strongest hippocampal activations in the late trials of each recording while cortical activations are stronger in the early trials.

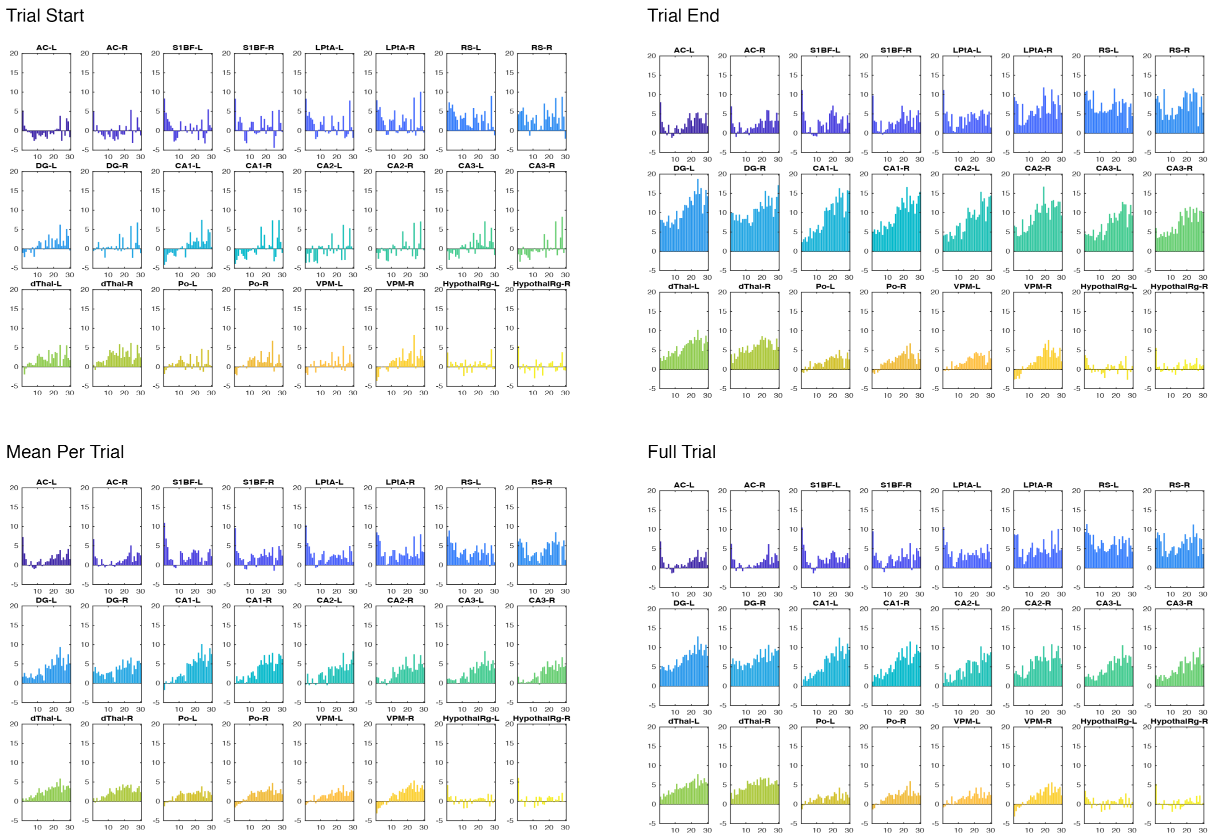


**Figure S6: Vascular reshaping affects the late component of single-trial hemodynamic responses more than its early component**

Different measures of hemodynamic reshaping grouped by timing from session start (0 refers to the first run in each recording) for 12 regions across individuals (N= 7 animals, 22 recordings). Left columns represent early runs while right columns represent late runs. We measured early CBV response (top left: trial start), late CBV response (top right: trial end), mean CBV response between start and end (bottom left: mean per trial), mean CBV responses over the full 12-second trial (bottom right: Full trial). Note that the hemodynamic reshaping effects are almost absent in the early part of the CBV response (trial start), except in the dorsal thalamus, while they are extremely prominent in the late part of the CBV response (trial end), especially the potentiating effect in the dorsal hippocampus. Mean responses (bottom left and right) provide a smoothed measure of hemodynamic reshaping which allow for a finer linear regression analysis.
